# Mitochondrial proteostatic stress disrupts mitoribosome biogenesis and translation

**DOI:** 10.1101/2025.09.05.673478

**Authors:** Hauke Holthusen, Victoria A. Trinkaus, Carina Fernandez Gonzalez, Itika Saha, Isabelle Pachmayr, Orsolya Kimbu Wade, Anja Deiser, Barbara Hummel, Patricia Yuste-Checa, Kenneth Ehses, Roman Körner, Rubén Fernández-Busnadiego, Ritwick Sawarkar, Ralf Jungmann, Mark S. Hipp, F. Ulrich Hartl

## Abstract

Protein aggregation in various cellular compartments is a hallmark of proteostasis impairment linked to aging and numerous pathologies. Mitochondrial function depends on a balanced interplay of proteins imported from the cytosol as well as those synthesized on mitochondrial ribosomes (mitoribosomes). Here, we reveal an unexpected susceptibility of mitoribosome biogenesis to organellar proteostatic stress. Importing aggregation-prone proteins into yeast and human mitochondria triggered a chain of detrimental events involving extensive co-aggregation of newly-imported mitoribosome subunits and other RNA-binding proteins, as well as local disruption of mitochondrial cristae morphology. As a result, mitoribosome assembly and mitochondrial translation were severely impaired, leading to respiratory deficiency and, ultimately, loss of mitochondrial DNA. Surprisingly, dysfunction of mitochondrial HSP60 phenocopied the ribosome biogenesis defect and inhibition of translation, indicating a pronounced chaperone dependence of mitoribosome proteins. Declining mitochondrial translation likely contributes to aging and diseases associated with deficiencies in mitochondrial protein quality control machinery.

## INTRODUCTION

Cells invest extensively in protein quality control machinery to maintain protein homeostasis (proteostasis) in the face of external and internal stress conditions.^1–3^ When these protective mechanisms fail or are overtaxed, as is increasingly the case during aging, toxic protein aggregates accumulate – a process that is linked to a wide range of diseases.^1–3^ The consequences of protein aggregation have been studied primarily in the cytosol, the main site of protein synthesis. Less is known about the mechanisms of aggregate toxicity in the nucleus, endoplasmic reticulum (ER) and mitochondria.^3,4^ Understanding the vulnerability of mitochondria to this form of stress is of particular interest, as a decline in mitochondrial function is associated with normal aging and age-related pathologies.^5,6^ Notably, mitochondria represent a major compartment of protein folding and assembly, both for proteins imported from the cytosol and synthesized within the organelles.^7^ Moreover, since mitochondria are of endosymbiotic bacterial origin, the limitations and vulnerability of their protein quality control machinery may differ from other cellular compartments.

Mitochondria are essential eukaryotic organelles of metabolism and energy production. Features reflecting their endosymbiotic ancestry include a double membrane architecture, a minimal circular genome, and a complete system for transcription and protein translation.^8^ The mitochondrial DNA (mtDNA) is organized into numerous protein-DNA complexes called nucleoids.^9^ mtDNA copy number per cell is used as a proxy for mitochondrial health and is reduced in aged individuals, mitochondrial diseases and age-related neurodegenerative disorders.^10,11^

While only a small number of proteins (8 in *S. cerevisiae* and 13 in humans) are synthesized within the organelles, ∼1000 different mitochondrial proteins are synthesized in the cytosol and must be imported into one of the mitochondrial subcompartments: outer membrane, inner membrane, intermembrane space and matrix.^7,12–14^ Proteins of the matrix traverse the translocation machinery of the outer and inner membranes in an unfolded state, followed by chaperone-assisted folding.^15^ The biogenesis of the respiratory complexes of the inner membrane requires the coordinated production and assembly of hydrophobic membrane proteins made inside the organelle and subunits imported from the cytosol,^16–18^ with the risk of producing unassembled subunits and assembly intermediates when this coordination fails.^19–21^ The conformational integrity of mitochondrial proteins is further challenged by the high levels of reactive oxygen species generated as a byproduct of respiration.^22,23^ Proteins from cytosolic aggregates may enter mitochondria and additionally burden matrix proteostasis.^24^ Protein misfolding and aggregation in mitochondria has been observed under a variety of conditions,^25–27^ with mitochondrial chaperones and proteases being upregulated via the mitochondrial unfolded protein response (mtUPR).^28–30^ However, the dominant mitochondrial stress response in mammalian cells is the integrated stress response (ISR),^31–33^ characterized by cytoplasmic translational attenuation through eIF2α phosphorylation^34,35^ and upregulation of the transcription factor ATF4 to activate chaperones and anti-oxidative proteins.^36^

Folding and assembly of proteins imported into the mitochondrial matrix is mediated by the essential chaperones mtHSP70 and HSP60/HSP10, which are closely related to bacterial DnaK and GroEL/GroES, respectively.^15^ Terminally misfolded proteins or excess unassembled subunits of multiprotein complexes are degraded by the bacterial-type matrix proteases LONP1 and CLPX/P or by the membrane proteases i-AAA and m-AAA.^37^ Mutations in HSP60 are associated with the neurodegenerative disorders spastic paraplegia 13 (SPG13) and mitCHAP-60 disease, while mutations in LONP1 are associated with the multisystem developmental disorder CODAS syndrome.^38–40^ The global decline in chaperone capacity during aging may also compromise the mitochondrial proteostasis system.^2,3^

To systematically analyze the consequences of proteostatic stress in mitochondria, we targeted aggregation-prone artificial β-sheet proteins^41,42^ to the mitochondrial matrix in *Saccharomyces cerevisiae* and human cells. Variants of these proteins have previously been used to explore gain-of-function toxicity of aggregation in the cytosol, nucleus and ER.^42–44^ Alternatively, we imported mutationally destabilized mitochondrial ornithine transcarbamylase (OTCΔ).^30^ Both model proteins formed similar aggregate foci within the mitochondrial matrix, disrupted cristae morphology, and caused pronounced growth impairment under respiratory conditions. As a primary insult, the mitochondrial aggregates prominently sequestered newly-imported mitoribosome proteins (mtRPs) and other RNA-binding proteins (mtRNABP). This resulted in defective ribosome assembly and severe impairment of mitochondrial translation with reduced abundance of respiratory chain complexes, and ultimately, mtDNA loss. Interestingly, the distinct conformational vulnerability of mtRPs correlated with their pronounced dependency on HSP60 for folding. Loss of HSP60 function resulted in degradation of mtRPs by LONP1 and in extensive aggregation when both HSP60 and LONP1 were defective. The nuclear-encoded HSP60/LONP1 axis thus controls the level of functional mitoribosomes. These results elucidate the mechanisms underlying the toxicity of protein aggregation in mitochondria and have relevance in aging and diseases associated with mitochondrial proteostatic stress.

## RESULTS

### Mitochondrial protein aggregates impair respiratory function

To generate protein aggregates in mitochondria of *S. cerevisiae* and human cells, we used an artificial β-sheet protein (β23) that was designed to form amyloid-like fibrils.^42–45^ This 63 amino acid protein consists of 6 β-strands connected by short linker sequences, with each β-strand comprising 7 alternating polar and non-polar residues (Figure 1A).^45^ β23 has no evolved biological function, but similar bipolar sequences occur in ∼30% of human proteins. For mitochondrial targeting, we fused the cleavable mitochondrial targeting sequence (MTS) of yeast Fo-ATP synthase subunit 9 to the N-terminus of β23 (hereafter referred to as Mito-β) (Figure 1A). A myc epitope-tag was added for detection. As an additional model protein, we chose an unstable version of rat mitochondrial ornithine transcarbamylase truncated from amino acid 30 to 114 (OTCΔ) (Figure 1A), previously used to analyze mitochondrial protein aggregation.^30,46^ Mitochondria-targeted myc-GFP (Mito-GFP) served as a non-aggregating control protein.

**Figure 1.**
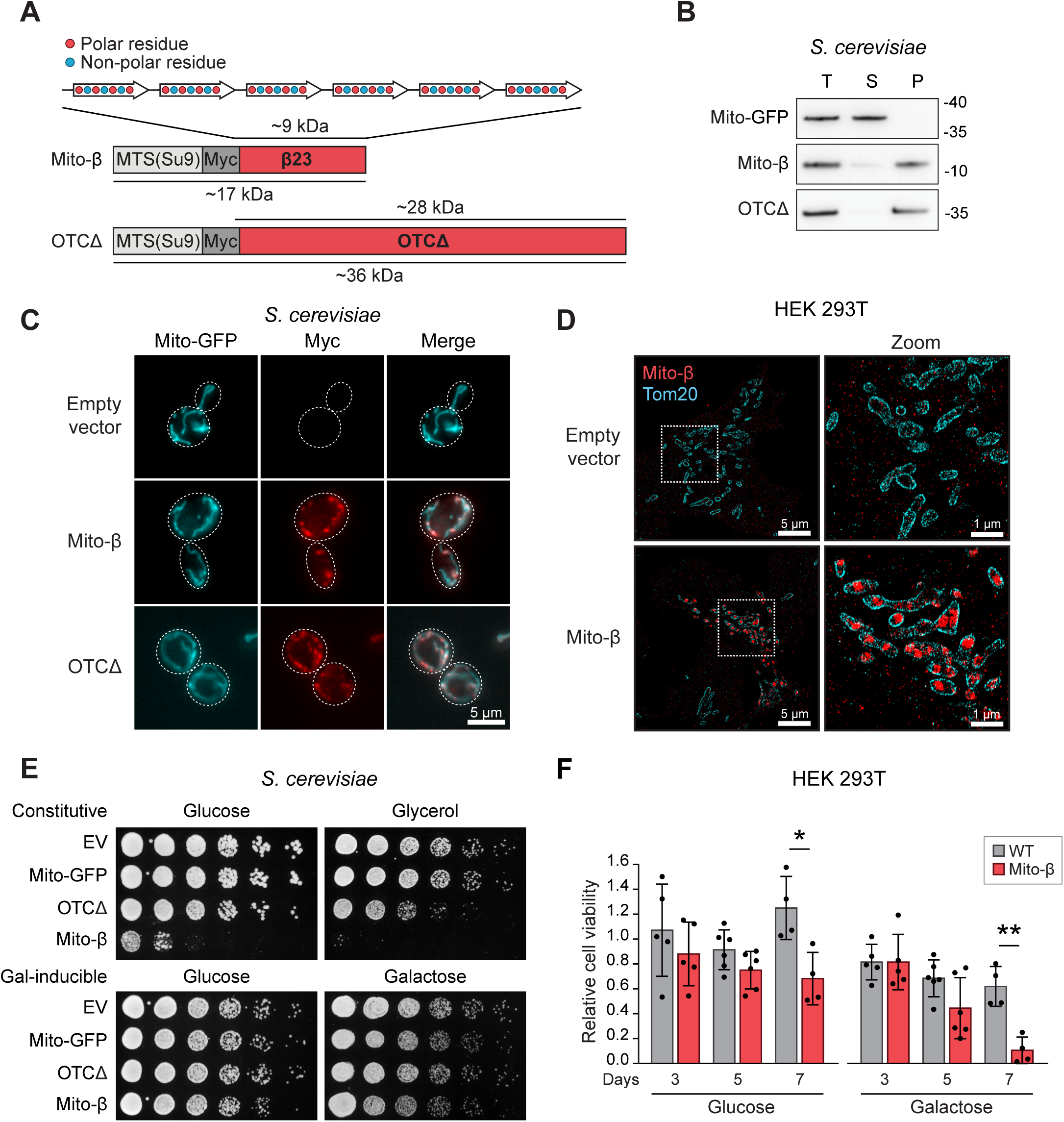
Mitochondrial protein aggregates impair respiratory function. (A) Schematic depiction of Mito-β and OTCΔ. MTS(Su9), mitochondrial targeting sequence of ATP-synthase subunit 9 from *Neurospora crassa*; myc, myc epitope tag; β-sheet protein, model protein designed to form β-sheet containing aggregates; OTCΔ, truncated ornithine transcarbamylase, aa 30-114 deleted. (B) Insolubility assay of galactose-induced Mito-GFP, Mito-β and OTCΔ. Yeast cells were lysed in buffer containing 1% Triton X-100 and lysates were fractionated by centrifugation, followed by immunoblotting with anti-myc antibody. T: total, S: soluble, P: pellet. (C) Immunofluorescence staining of Mito-β and OTCΔ in yeast cells co-expressing Mito-GFP. Constitutively expressed Mito-β and OTCΔ were detected with anti-myc antibody (red) and the mitochondrial matrix was visualized by Mito-GFP fluorescence (cyan). (D) Immunofluorescence staining of Mito-β in HEK 293T cells by DNA-PAINT super-resolution microscopy. Mito-β was detected with anti-myc antibody (red) and the mitochondrial outer membrane protein TOM20 by anti-TOM20 antibody (cyan). (E) Spot dilution assays of yeast cells with constitutive or galactose-induced expression of Mito-GFP, Mito-β or OTCΔ in glucose, galactose or glycerol medium. (F) Determination of cellular viability of HEK 293T cells by MTT-assay. Wild type (WT) cells and cells expressing Mito-β under a doxycycline (Dox)-inducible promoter were cultured for up to 7 days in glucose or galactose containing medium with and without 0.5 µg/mL Dox. Viability of +Dox cells is displayed relative to -Dox cells. Mean with error bars as SD (n ≥ 4). p-Values by One-way ANOVA with Bonferroni′s multiple comparisons test. *p < 0.05, **p < 0.01.

We expressed these proteins in yeast cells under the constitutive *GPD* promoter or under the inducible *GAL1* promoter. The targeting sequences of all three proteins were efficiently cleaved, as demonstrated by immunoblotting (Figure S1A). Only a small fraction of the total Mito-β remained in precursor form, indicating that targeting to the mitochondrial matrix occurred with high efficiency (Figure S1A). While Mito-GFP was soluble, Mito-β and OTCΔ were almost completely recovered in the insoluble fraction upon cell fractionation (Figure 1B), indicative of aggregation.

Immunofluorescence analysis showed that Mito-β and OTCΔ formed multiple distinct foci that co-localized with the mitochondrial network visualized by co-expression of Mito-GFP (Figure 1C). There was no apparent disruption of network morphology. Similar results were obtained when Mito-β and OTCΔ were expressed under the CMV promoter in HEK 293T cells 24 h after transfection of the constructs. Both proteins were again recovered mainly in the detergent-insoluble fraction (Figure S1B). While Mito-β was efficiently processed, some OTCΔ accumulated as the precursor form (Figure S1B). We visualized Mito-β in HEK 293T cells by DNA-PAINT super-resolution microscopy with TOM20 as a marker of the outer mitochondrial membrane, revealing the presence of large (up to ∼300 nm) aggregate foci (Figure 1D). These inclusions occupied areas of the mitochondrial matrix that were nucleoid free, as shown by DNA antibody staining (Figure S1C).

We next tested whether the mitochondrial aggregates caused cytotoxic effects. Yeast cells expressing Mito-β or OTCΔ were grown on fermentative (glucose) and respiratory (glycerol) plates. In glucose medium, yeast cells generate ATP mainly by glycolysis, whereas energy generation in glycerol medium is strictly dependent on mitochondrial respiration.^47,48^ Constitutive expression of Mito-β strongly inhibited growth on both glucose and glycerol medium, whereas OTCΔ reduced growth on glycerol medium only, suggesting a lower degree of toxicity (Figure 1E). Mito-GFP did not cause any growth impairment (Figure 1E). Transcriptome analysis showed that Mito-β-expressing cells were defective in the upregulation of nuclear-encoded subunits of respiratory chain complexes upon diauxic shift from glucose to glycerol medium (Figure S1D). Concomitantly, the transcriptional upregulation of chaperones of the mitochondrial matrix and cytosol, normally observed upon diauxic shift,^49–51^ was markedly enhanced (Figure S1E), indicative of general stress response activation in Mito-β-expressing cells. Constitutive expression of Mito-β increased the levels of reactive oxygen species (ROS) in glucose and glycerol medium by ∼2.4- and ∼1.5-fold, respectively (Figure S1F). Notably, induction of Mito-β from the *GAL1* promoter, although resulting in higher expression, was only mildly toxic (Figures 1E and S1A), for reasons that will become clear below.

Long-term expression of Mito-β in HEK 293T cells under doxycycline control also strongly impaired cell viability under conditions of high respiratory dependency (galactose medium),^52^ while toxicity was less pronounced under conditions of low respiratory dependency (glucose medium) (Figure 1F). Expression of Mito-β for 24 h activated the ISR with an increase in p-eIF2α comparable to that after 5 h treatment with the strong ISR activator tunicamycin (Figure S1G).

In summary, accumulation of protein aggregates in mitochondria inhibits cell growth in both yeast and human cells, especially under respiratory conditions, and results in stress response activation.

### Protein aggregates impair mitochondrial cristae architecture

To visualize the aggregates and their possible effects on mitochondrial architecture, we analyzed cells by cryo-electron tomography (cryo-ET). Yeast cells with galactose-induced expression of Mito-β or OTCΔ were grown without overt toxicity (Figure 1E). 3D reconstruction of tomograms showed that their mitochondria contained large aggregate inclusions, apparently amorphous in structure, occupying most of the volume of the matrix compartment (Figure S2). Mitochondria in control cells were elongated with cristae protruding into the matrix. In contrast, mitochondria with aggregates showed abnormal cristae structures. Furthermore, upon shift from glucose to glycerol medium for 24 h with constitutive expression of Mito-β, aggregate-containing mitochondria were swollen and completely lacking cristae, consistent with defective mitochondrial respiration and growth impairment (Figure S2).

Cryo-ET of HEK 293T cells also revealed Mito-β aggregates occupying much of the matrix volume of the mitochondria (Figure 2A). The aggregates appeared more reticular and less dense than in yeast cells. Again, cristae morphology was strongly disrupted (Figure 2A). To visualize the effect on cristae in the context of the mitochondrial network, we performed structured illumination microscopy on live HeLa cells, in which mitochondria are more interconnected than in HEK 293T cells (Figures 1D and 2B). We expressed Mito-β as a fusion protein with mScarlet (Mito-mScarlet-β) and stained cristae with MitoBright Green, a dye that accumulates in the mitochondrial intermembrane space and cristae lumen.^53^ Cristae were absent in areas of the mitochondrial network where Mito-mScarlet-β aggregates accumulated, but were largely unaffected in other areas (Figure 2B). Thus, the aggregate inclusions locally disrupt the mitochondrial architecture by creating cristae-free areas around them.

**Figure 2.**
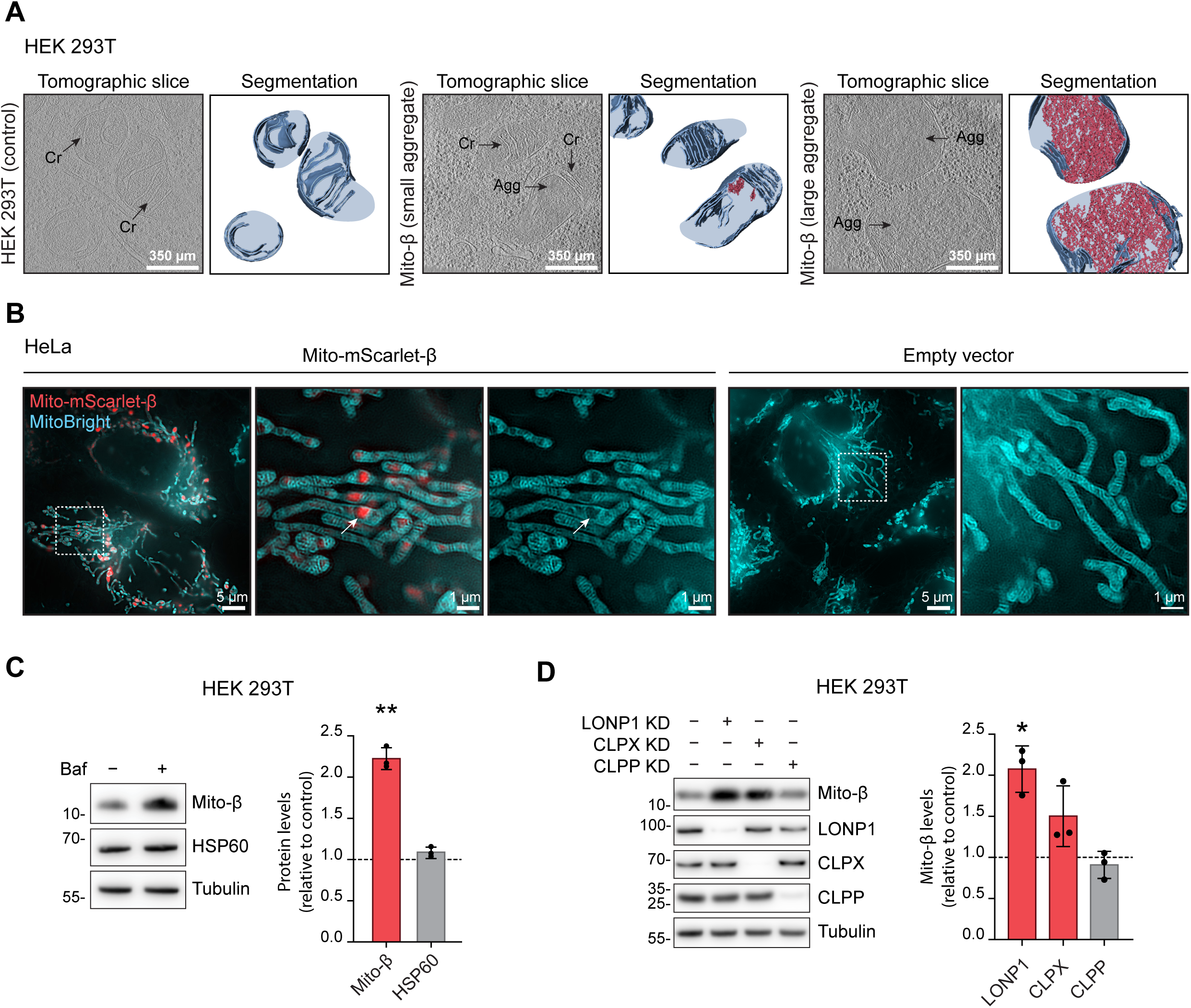
Protein aggregates impair mitochondrial cristae architecture. (A) Cryo-electron tomographic slices (1.4 nm thickness) of HEK 293T cells expressing Mito-β. WT cells and cells containing Dox-inducible Mito-β were cultured for 3 days with Dox. Segmentations show mitochondrial membranes including cristae (Cr) in blue and aggregates (Agg) in red. Tomograms were acquired from at least 3 different cells from two biological replicates. Scale bar: 350 µm. (B) Structured illumination microscopy of cristae in live HeLa cells transiently expressing Mito-mScarlet-β for 3 days. Cristae were stained with MitoBright green dye (cyan) and Mito-β was visualized by the fluorescence of mScarlet (red). Cells transfected with empty vector (EV) were used as control. Dotted square indicates magnified region on the right. White arrows highlight a Mito-β aggregate in a cristae-free area. (C) Quantification of Mito-β by immunoblotting after inhibition of autophagy in HEK 293T cells. Mito-β was induced with Dox for 3 days before cells were treated with 25 nM bafilomycin (Baf) for 24 h. Quantification of Mito-β and HSP60 levels in Baf treated cells is shown as bar graph relative to untreated cells. Mean with error bars as SD (n = 3). p-Values by Repeated-measures One-way ANOVA with Bonferroni′s multiple comparisons test. **p < 0.01. (D) Quantification of Mito-β in cells with knockdown (KD) of LONP1, CLPX and CLPP by immunoblotting with antibodies as indicated. Mito-β was induced with Dox for 4 days. Cells were transfected with siRNA for 4 days. Quantification of Mito-β levels is shown as bar graph relative to non-targeted siRNA control. Mean with error bars as SD (n = 3). p-Values by Repeated-measures One-way ANOVA with Dunnett′s multiple comparisons test. *p < 0.05.

Mitochondrial aggregates can induce mitophagy for selective degradation.^46,54,55^ Thus areas devoid of cristae may be targeted by mitophagy for degradation of Mito-β aggregates in lysosomes. Indeed, we observed that treatment of Mito-β-expressing HEK 293T cells with the lysosomal acidification inhibitor bafilomycin for 24 h resulted in a ∼2.2-fold increase in Mito-β levels, while HSP60 levels remained unchanged (Figure 2C), suggesting selective lysosomal degradation of Mito-β aggregates.

To determine whether mitochondrial matrix proteases are also involved in the degradation of Mito-β aggregates, we used RNA interference to downregulate LONP1, CLPX or CLPP by ∼85%. Downregulation of LONP1 increased Mito-β levels by ∼100% and downregulation of CLPX by ∼50%, while reducing CLPP had no effect (Figure 2D). CLPX is the unfoldase of the CLPX/P protease complex, but also functions independently of CLPP.^37,56^

In conclusion, Mito-β aggregation locally disrupts mitochondrial cristae structure. Mito-β degradation is mediated by the lysosome and by LONP1, possibly in cooperation with CLPX.

### Mitochondrial aggregates sequester mtRPs

Protein aggregates may cause cytotoxic effects by interacting with endogenous proteins, resulting in their sequestration and (partial) loss of function.^3,57,58^ To test whether the mitochondrial aggregates exert toxicity by a similar mechanism, we performed an interactome analysis of Mito-β and OTCΔ aggregates by quantitative mass spectrometry (LC-MS/MS) (Figures 3A and S3A-C; Table S1A). Mito-β aggregates were immunoprecipitated from yeast lysates after 24 h of galactose-induced expression. Cells transformed with empty vector (EV) or Mito-GFP were used as controls. Mito-β interactors were almost exclusively mitochondrial proteins. Gene ontology (GO)-term analysis (Figure 3A) revealed a significant enrichment of several functional groups of proteins involved in mitochondrial gene expression (Figure 3A), including mitoribosome proteins (mtRPs) (50 of the 80 imported mtRPs and the mt-encoded Var1) (Figure S3A), mitochondrial nucleoid and RNA-binding proteins (mtRNABPs) (Figures S3A and S3B), and several mitochondrial chaperones and other protein quality control factors (Hsp60, mtHsp70 [Ssc1], Mdj1, Hsp78, LonP1 [Pim1], ClpX) (Figure S3C). In addition, proteins involved in iron-sulfur-cluster (ISC) biogenesis (Ssq1, Isu1/2, Isu1/2 and Nfu1) and ISC-containing proteins (Lip5, Rip1, Ilv3, Hem15, Aco1, Sdh2 and Lys4), as well as most TCA cycle enzymes and ATP-synthase subunits were enriched in the aggregate interactome (Figure S3C). Notably, the interactome of OTCΔ aggregates, although less complex, showed almost complete overlap with the Mito-β interactome, again including numerous mtRPs (Figure 3B; Table S1B). Thus, the mitochondrial aggregates formed by different proteins have very similar interaction properties.

**Figure 3.**
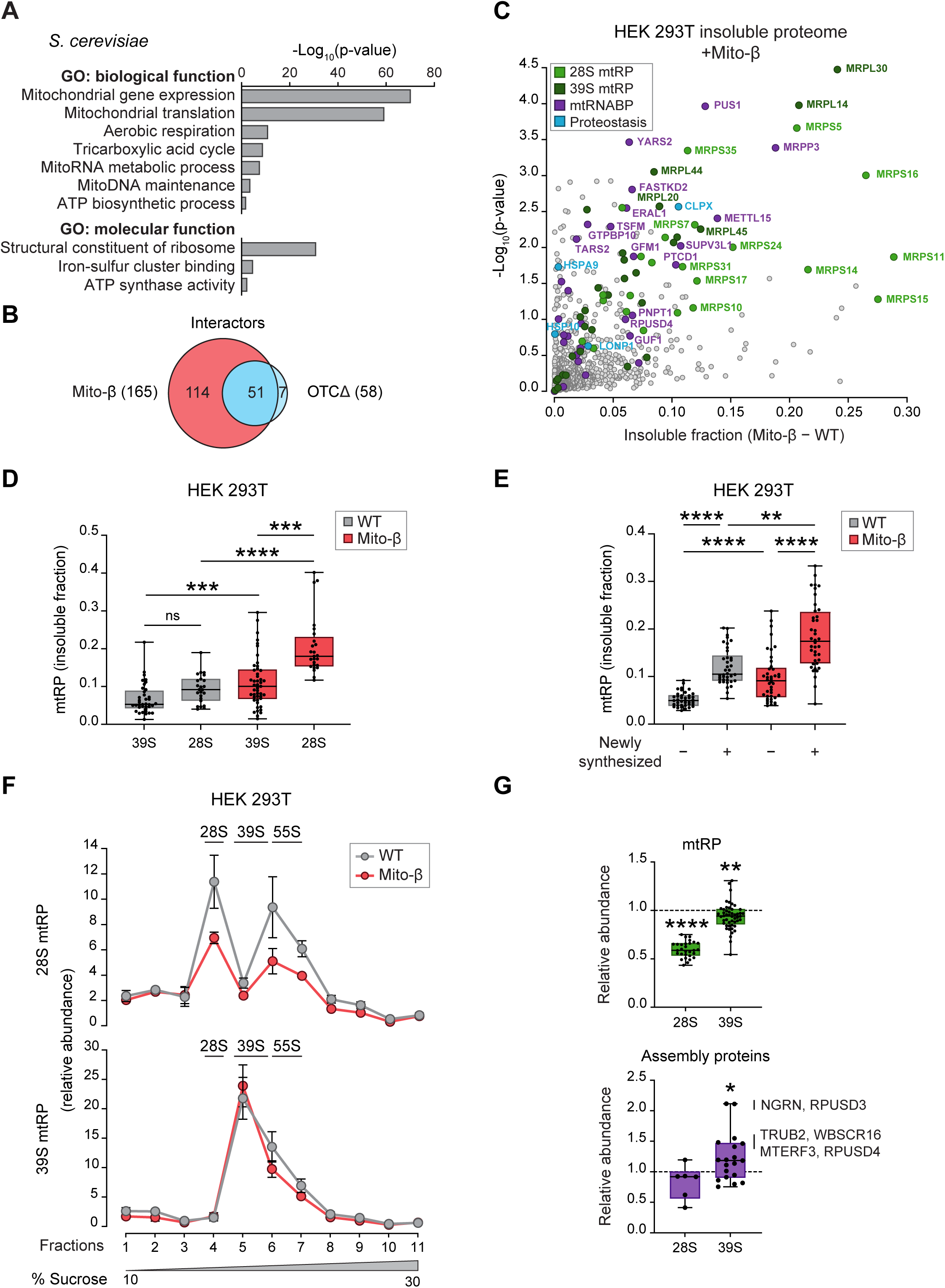
Mitochondrial aggregates sequester mtRPs. (A) Interactome analysis of Mito-β in yeast cells by SILAC LC-MS/MS displayed as GO-term enrichment of proteins with at least 4-fold enrichment over Mito-GFP. Selected GO-terms are shown. See also Figures S3A, S3B and S3C and Table S1. (B) Overlap between interactors of Mito-β and OTCΔ aggregates. Only proteins with at least 2-fold enrichment over the EV control (p < 0.05) were included. Numbers of proteins are indicated. See also Table S1. (C) Insoluble proteome analysis of Mito-β in HEK 293T by label-free LC-MS/MS. p-Value is plotted against insoluble fraction (pellet/total, from LFQ-values). Background insoluble fraction from wild-type (WT) control cells was subtracted. WT cells and cells containing Dox-inducible Mito-β were cultured for 3 days with Dox. Selected groups of proteins are highlighted. See also Table S1. (D) Analysis of the insoluble fraction of mtRPs in (C) according to 39S and 28S subunit proteins. Boxes and whiskers indicate quartiles and the line indicates the median (n=4). p-Values by Mixed-effects analysis with Bonferroni′s multiple comparisons test. ***p < 0.001, ****p < 0.0001. (E) Analysis of insoluble fraction of newly-imported mtRPs. Mito-β expression was induced for 16 h in HEK 293T before cells were grown in heavy isotope for 4 h to label newly-translated proteins. Insoluble proteome was analyzed by SILAC LC-MS/MS. Box plot shows insoluble fraction (pellet/total) of mtRPs in HEK 293T cells with and without Mito-β expression. Boxes and whiskers indicate quartiles and the line indicates the median (n = 5). p-Values by Two-way ANOVA with Tukey’s multiple comparison test. **p < 0.01, ****p < 0.0001. See also Table S1. (F-I) Analysis of mitoribosome assembly by sucrose gradient fractionation of Mito-β-expressing HEK 293T cells, followed by label-free LC-MS/MS. Average abundance (summed LFQ-values) of 28S and 39S mtRPs was plotted per fraction (F). Boxplots (I) show abundance of 28S and 39S mitoribosome and assembly proteins in fractions corresponding to assembled mitoribosomes of Mito-β-expressing HEK 293T cells relative to WT (summed LFQ values from fraction 4-7 for 28S and fraction 5-7 for 39S). Specific assembly proteins of the pseudouridinylation module are indicated. Boxes and whiskers indicate quartiles and the line indicates the median. (n=3). p-Values by Mixed-effects analysis with Bonferroni′s multiple comparisons test. *p < 0.05, **p < 0.01, ****p < 0.0001. See also Table S2.

To determine the extent to which specific proteins were depleted from the soluble pool by co-aggregation with Mito-β, we analyzed the detergent-insoluble proteome of Mito-β-expressing yeast cells in comparison to control cells containing empty vector (EV). Strikingly, numerous mtRPs were highly insoluble (Figure S3D; Table S1C), with the insoluble fraction increasing from 5-15% to 30-50% of the total (Figure S3E). mtRNABPs, proteostasis components, and ISC proteins were also partially insoluble upon expression of Mito-β, whereas TCA cycle components and ATP-synthase subunits were only slightly affected (Figure S3E).

Insoluble proteome analysis of Mito-β aggregates in HEK 293T cells reproduced the results obtained in yeast. mtRPs and mtRNABPs were rendered strongly insoluble in the presence of the aggregates, with the insoluble fraction of some proteins exceeding 25% of the total (Figure 3C; Table S1D). Insolubility was most pronounced for proteins of the small mitoribosomal subunit (28S) (Figures 3C, 3D and S3F) and varied considerably between individual proteins, suggesting that co-aggregation with Mito-β occurred prior to ribosome assembly. Interestingly, several of the most insoluble mtRNABPs are involved in mitoribosome assembly and rRNA processing (ERAL1, METTL15, FASTKD2, PTCD1, MRPP3).

To determine whether the aggregates preferentially sequester newly-imported mitochondrial proteins, we analyzed the insoluble fraction of Mito-β-expressing HEK 293T cells after labeling with amino acid isotopes. Mito-β was induced for 16 h to accumulate aggregates, followed by labeling of newly-imported proteins with heavy lysine and arginine for 4 h. Notably, in wild type (WT) cells, the insoluble fraction of newly-imported mtRPs was higher than that of their preexisting counterparts (Figure 3E; Table S1E), suggesting that normally a fraction (∼10-15%) of imported mtRPs fail to fold or assemble. The insolubility of newly-imported mtRPs was substantially increased in the presence of Mito-β, indicating that the mtRPs are captured by the aggregates preferentially prior to ribosome assembly.

We next investigated whether ribosome assembly is disrupted in aggregate containing mitochondria. Mitochondrial lysates from HEK 293T cells were separated by sucrose gradient fractionation, followed by MS analysis. We found that mtRPs of the small subunit were reduced by ∼40% on average in the 28S and 55S mitoribosome fractions (Figures 3F and 3G; Table S2A), consistent with the preferential insolubility of the 28S subunit proteins (Figures 3C and 3D). While the gradient fractionation did not allow a clear separation of 39S and 55S ribosomes, the overall abundance of the assembled 39S mtRPs appeared unaffected (Figures 3F and 3G). However, mtRNABPs belonging to the pseudouridinylation module of the 39S subunit rRNA were enriched in 39S fractions from aggregate containing mitochondria,^59,60^ suggesting an accumulation of late-stage 39S assembly intermediates (Figure 3G). The levels of mt-encoded 12S and 16S rRNA, essential components of 28S and 39S ribosomes, respectively, were normal as determined by qPCR, suggesting that rRNA transcription was not impaired in the presence of Mito-β aggregates (Figure S3G).

In conclusion, protein aggregation in mitochondria results in extensive co-aggregation and sequestration of numerous newly-imported mtRPs and mtRNABPs, manifesting in defective mitoribosome assembly.

### Mitochondrial aggregates impair mitochondrial translation

We next investigated whether the defect in mitoribosome assembly induced by the aggregates impaired mitochondrial protein synthesis. We first tested this in yeast cells using a reporter strain engineered to express mt-encoded GFP.^61^ Mito-β or OTCΔ expression induced with galactose for 24 h resulted in a significant reduction in GFP levels by 30-40%, while the abundance of nuclear-encoded mitochondrial Hsp60 remained unchanged (Figure 4A). Consistent with reduced mitochondrial protein synthesis, the level of mt-encoded subunit 2 of cytochrome oxidase (Cox2) was reduced by 50-60% (Figure 4A). Interestingly, the level of cytochrome c oxidase subunit 4 (Cox4), which is nuclear encoded and thus imported into the mitochondria, was reduced to a similar extent (Figure 4A), suggesting that cytochrome oxidase (complex IV) assembly was broadly affected. This finding prompted us to analyze the total proteome of yeast cells expressing Mito-β (Figure 4B). We quantified 568 nuclear-encoded mitochondrial proteins and 4 mt-encoded proteins by label-free MS (Table S3). While the nuclear-encoded proteins were not globally reduced, expression of Mito-β resulted in a reduction in the levels of multiple imported respiratory chain subunits (Figures 4B and S4A-B), suggesting that unassembled proteins were degraded in the absence of mt-encoded partner subunits.^62,63^ Notably, the overall abundance of mtRPs was not reduced, but rather slightly increased. A possible explanation for this is that the high degree of insolubility of these proteins due to aggregation prevented their degradation (Figures 4B and S4B).

**Figure 4.**
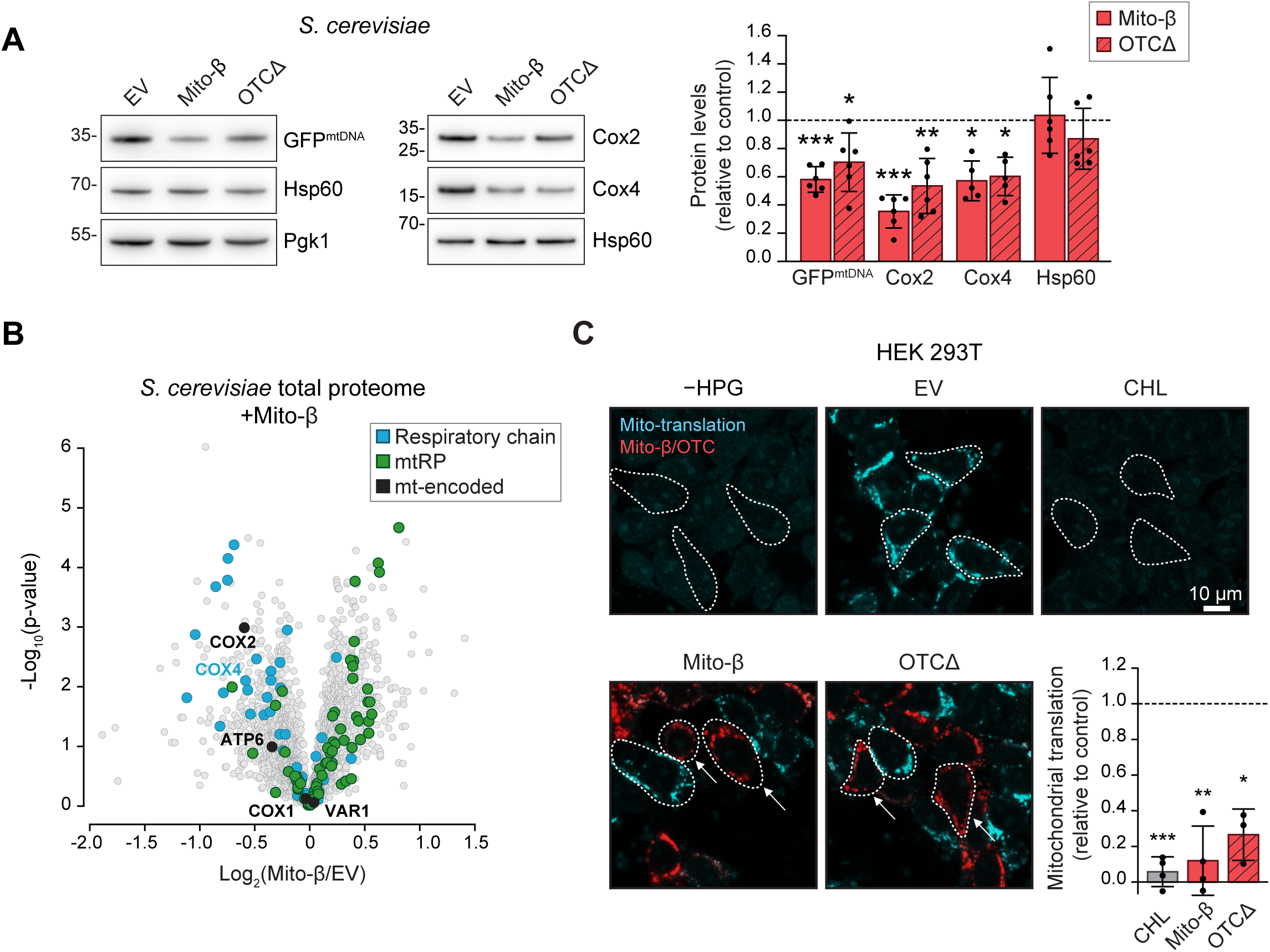
Mitochondrial aggregates impair mitochondrial translation. (A) Analysis of mitochondrial translation efficiency in yeast cells with mitochondrial aggregates. Levels of mt-encoded GFP (GFP^mtDNA^), cytochrome-c oxidase subunits 2 (COX2; mt-encoded) and 4 (COX4; nuclear-encoded) were analyzed by immunoblotting. EV control cells and cells expressing galactose-inducible Mito-β or OTCΔ were grown in galactose medium for 24 h. Quantification of protein amounts by densitometry is shown as bar graph. Mean with error bars as SD. p-Values by Repeated-measures One-way ANOVA with Dunnett′s multiple comparisons test (GFP^mtDNA^) and Mixed-effects analysis with Bonferroni′s multiple comparisons test (COX2/4). GFP n = 6, COX2 n = 5, HSP60 n = 5, COX4 n = 4. *p < 0.05, **p < 0.01, ***p < 0.001. (B) Analysis of total proteome of Mito-β-expressing yeast cells by label-free LC-MS/MS. Volcano plot of Mito-β against EV control (fold change of LFQ-values). Cells were grown for 24 h in galactose medium. See also Table S3. (C) Mitochondrial translation in HEK 293T cells containing mitochondrial aggregates. Newly-synthesized mitochondrial proteins were labeled with L-homopropargylglycine (HPG) and incorporation detected by Click chemistry (cyan). Chloramphenicol (CHL) (100 µg/µL) treated cells served as positive control. Mito-β and OTCΔ transfected cells were visualized by immunofluorescence (red) and selected for quantification of mean fluorescence intensity (10 cells per repeat). Translation in control cells is set to 1 (dotted line). Mean with error bars as SD. p-Values by Mixed-effects analysis with Dunnett′s multiple comparisons test. Mito-β and CHL (n = 4), OTCΔ (n = 3), *p < 0.05, **p < 0.01, ***p < 0.001.

To explore whether co-aggregation of mtRPs similarly leads to impairment of mitochondrial translation in HEK 293T cells, we selectively labeled mitochondrial translation products with the methionine analog homopropargylglycine (HPG), while inhibiting cytosolic translation with emetine. HPG contains an alkyne moiety that allows detection with a fluorescent azide by Click chemistry.^64^ Strikingly, HPG incorporation in cells containing Mito-β or OTCΔ aggregates was reduced by ∼90% and ∼70%, respectively, compared to control cells, similar to inhibition of mitochondrial translation with chloramphenicol (CHL) (Figure 4C). Thus, translation efficiency is reduced by a greater amount than suggested by the degree of mtRP insolubility alone (20-30%; Figure 3C), consistent with combinatorial effects on mtRPs and mtRNABPs, as well as defects in ribosome assembly. Indeed, the levels of mt-encoded COX2 protein were reduced by up to 70% after 7 days of Mito-β expression (Figure S4C), in line with the reduced viability of Mito-β- or OTCΔ-expressing HEK 293T cells in respiratory medium (Figure 1F).

A reduction in respiratory chain subunits due to protein aggregation in mitochondria was also observed in neuronal cells differentiated from human induced pluripotent stem cells (iNeurons). Mito-β and Mito-GFP were expressed by lentiviral transduction and mitochondrial localization was confirmed by immunofluorescence staining (Figure S4D). Mitochondrial aggregates formed in the soma and in neurites. The mt-encoded COX2 was reduced by ∼40% after 14 days of Mito-β expression, while HSP60 remained unchanged, suggesting reduced mitochondrial translation activity (Figure S4E). This reduction correlated with a ∼20% decrease in viability (Figure S4F).

Taken together, these results demonstrate that mitochondrial aggregates substantially impair mitochondrial gene expression and respiratory chain complex biogenesis, not only in yeast, but also in HEK 293T cells and human iNeurons.

### Mitochondrial aggregates cause mtDNA loss

mtDNA copy number and integrity decreases in an age-dependent manner in mice and humans,^10,11,65,66^ and during replicative aging in yeast,^67^ where maintenance of mtDNA depends on functional mitochondrial protein translation.^68,69^ Partial or complete loss of mtDNA in yeast cells results in respiratory deficiency and a small colony growth phenotype on fermentable carbon sources, termed petite.^70,71^ We investigated whether the formation of mitochondrial aggregates causes loss of mtDNA by measuring petite frequency as a proxy. Functional mitochondria in normal colonies reduce tetrazolium dye to formazan, which stains the colonies red, while respiratory-deficient petite colonies remain white.^72^ We observed a petite frequency of 83% in cells with Mito-β aggregates and a less pronounced but still highly significant petite frequency of 17% in cells with OTCΔ aggregates (constitutive Mito-β or OTCΔ expression), while petite occurrence in EV control cells was very low (∼1%) (Figure 5A).

**Figure 5.**
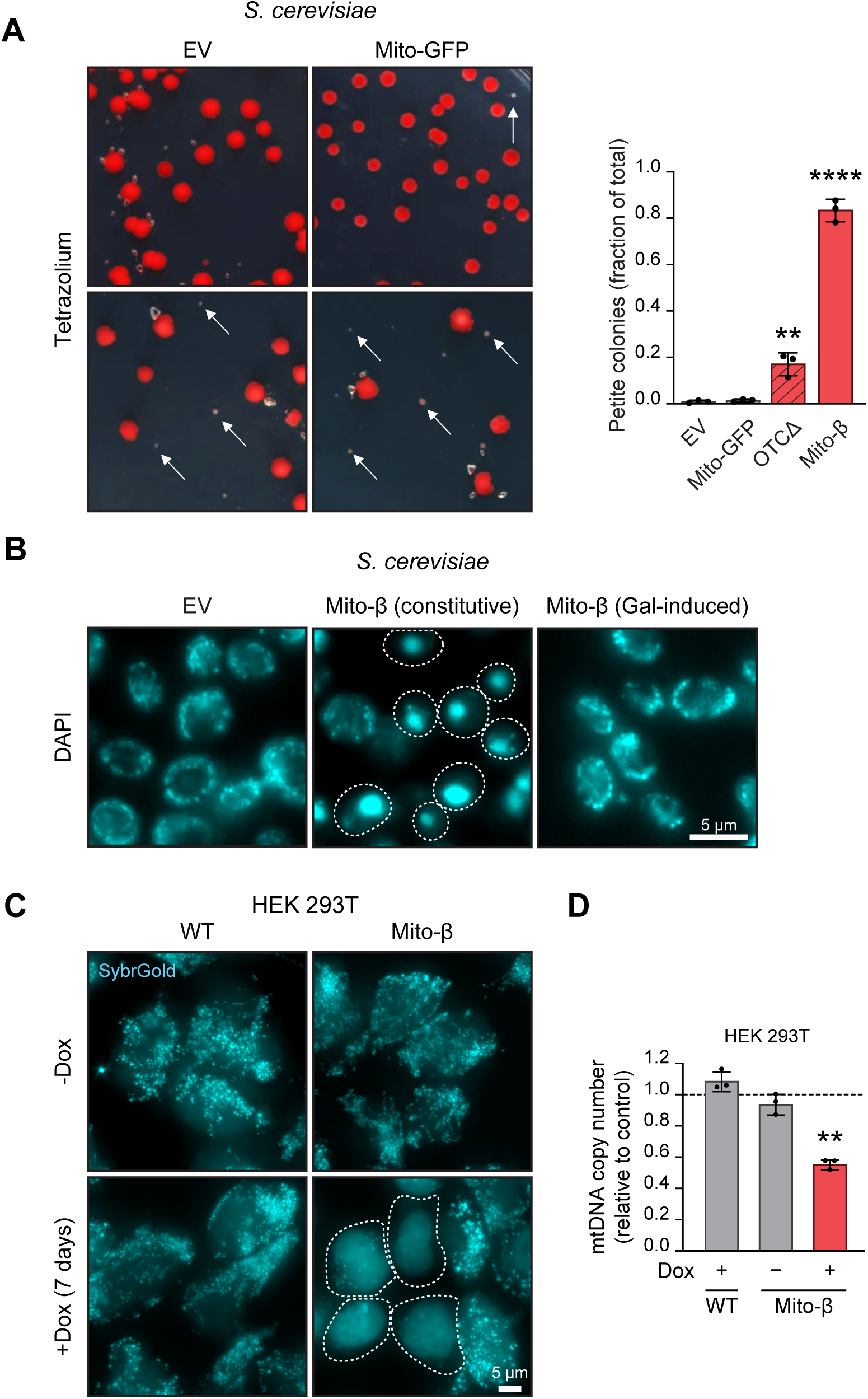
Mitochondrial aggregates cause mtDNA loss. (A) Quantification of petite colonies. Yeast cells constitutively expressing Mito-GFP, Mito-β or OTCΔ as well as EV control cells were grown on glucose plates and stained with Tetrazolium. White arrows indicate small, less colored colonies which represent the petite phenotype, in contrast to normal-sized, deep red colonies. Quantification of the fraction of petite colonies is shown as bar graph. Mean with error bars as SD (n = 3). p-Values by One-way ANOVA with Dunnett’s multiple comparison test. **p < 0.01, ****p < 0.0001. (B) Visualization of mtDNA in yeast cells. Cells expressing Mito-β either constitutively or upon induction with galactose for 24 h, and EV control cells were stained with DAPI and analyzed by live cell fluorescence microscopy. Cells were grown in galactose medium for 24 h. Boundaries of cells with minimal or absent mtDNA are outlined. (C) Visualization of mitochondrial DNA in HEK 293T cells. mtDNA in cells containing Dox-inducible Mito-β were cultured for 7 days with and without Dox and stained with SybrGold. WT cells were analyzed as control. Boundaries of cells with partial and severe loss of mtDNA are outlined. (D) Relative copy number of mtDNA in cells cultured as in (C), as quantified by qPCR. Mean with error bars as SD (n = 3). p-Values by Repeated-measures One-way ANOVA with Dunnett′s multiple comparisons test. **p < 0.01.

To verify that the petite phenotype was caused by loss of mtDNA, we visualized mtDNA using DAPI staining, which preferentially stains mtDNA in live yeast cells. Indeed, cells with constitutive Mito-β expression showed sparse mitochondrial DAPI staining consistent with mtDNA loss (Figure 5B). Notably, we observed a similar phenotype upon deletion of mtRPs, ATP-synthase subunits or mtDNA binding proteins (Figure S5A), proteins that interacted with the Mito-β aggregates (Figures S3A, S3B, S3C), consistent with reports that mtDNA maintenance depends on these factors.^73–76^ Thus, depletion of mitoribosomes would be sufficient to cause mtDNA loss. Of note, while DAPI stained predominantly the mitochondrial network in control cells, the nucleus became strongly DAPI positive upon mtDNA loss. This effect may be due to a higher concentration of DAPI in these cells, possibly as a result of increased membrane permeability. In contrast to constitutive Mito-β expression, galactose-induced expression of Mito-β for 24 h in liquid culture was not sufficient to cause mtDNA loss (Figure 5B), correlating with the absence of a growth defect (Figure 1E). However, after several generations on plates with galactose, Mito-β expression caused mtDNA loss and petite phenotype (Figure S5B). Thus, long-term expression of Mito-β results in high toxicity due to loss of mtDNA, explaining why short-term galactose-induced expression was less toxic compared to constitutive expression (Figure 1E).

Transcriptome analysis of yeast cells constitutively expressing Mito-β revealed stress signatures previously described for cells following mtDNA loss, including activation of the retrograde signaling response, pleiotropic drug response, and iron starvation response pathways (Figure S5C).^67,77–79^ Activation of the iron starvation response was accompanied by elevated iron levels in Mito-β-expressing cells, consistent with a defect in ISC assembly that signals cellular iron deficiency (Figure S5D).^80^ Of note, ISC deficiency may be related to sequestration of ISC biogenesis factors by Mito-β aggregates (Figure S3C).

Does mitochondrial protein aggregation also cause mtDNA loss in HEK 293T cells? To address this question, we expressed Mito-β in glucose medium for up to 7 days and visualized mtDNA by SybrGold fluorescence staining.^81^ After 3 days of Mito-β expression, 16% of cells had a low mtDNA count (<100 nucleoids), which increased to 21% after 5 days and 40% after 7 days, whereas only ∼7% of WT control cells had a low mtDNA count at all time points (Figures 5C and S5E). qPCR measurements confirmed the mtDNA decrease. After 7 days of Mito-β expression, mtDNA copy number was reduced by ∼45% compared to control cells (Figure 5D). Notably, the increase in the frequency of mtDNA loss between 5 and 7 days of Mito-β induction correlated with the impairment of cell viability observed at this time when cells were grown under respiratory conditions in galactose medium (Figure 1F). Accordingly, the loss of mtDNA as a consequence of protein aggregation in mitochondria strongly correlates with the onset of severe toxicity.

### Disruption of mitochondrial proteostasis machinery phenocopies effects of protein aggregates

The distinct propensity of mtRPs and mtRNABPs to be sequestered by protein aggregates suggested that these proteins may normally be dependent on molecular chaperones for folding and assembly. Mutations in the mitochondrial chaperonin HSP60 are known to cause the neurodegenerative diseases spastic paraplegia 13 (SPG13) and MitCHAP-60 disease.^38,39^ Strikingly, we found that a ∼70% knock-down (KD) of HSP60 in HEK 293T cells (Figure S6A) caused a ∼50% reduction in the levels of most mtRPs and numerous mtRNABPs, as well as the mt-encoded proteins COX2, COX3 and ATP8 (Figures 6A and 6B; Table S4A and S4B). MS analysis of sucrose gradient fractions confirmed a decrease in the level of assembled mitoribosomes, with 39S and 28S subunit complexes being reduced by ∼50% and ∼28%, respectively (Figures 6C and S6B; Table S2A). Ribosome assembly factors in ribosome fractions were reduced by ∼50% (Figure 6C). Indeed, mitochondrial translation was inhibited by ∼90%, as measured by HPG labeling (Figures 6D and S6C).

**Figure 6.**
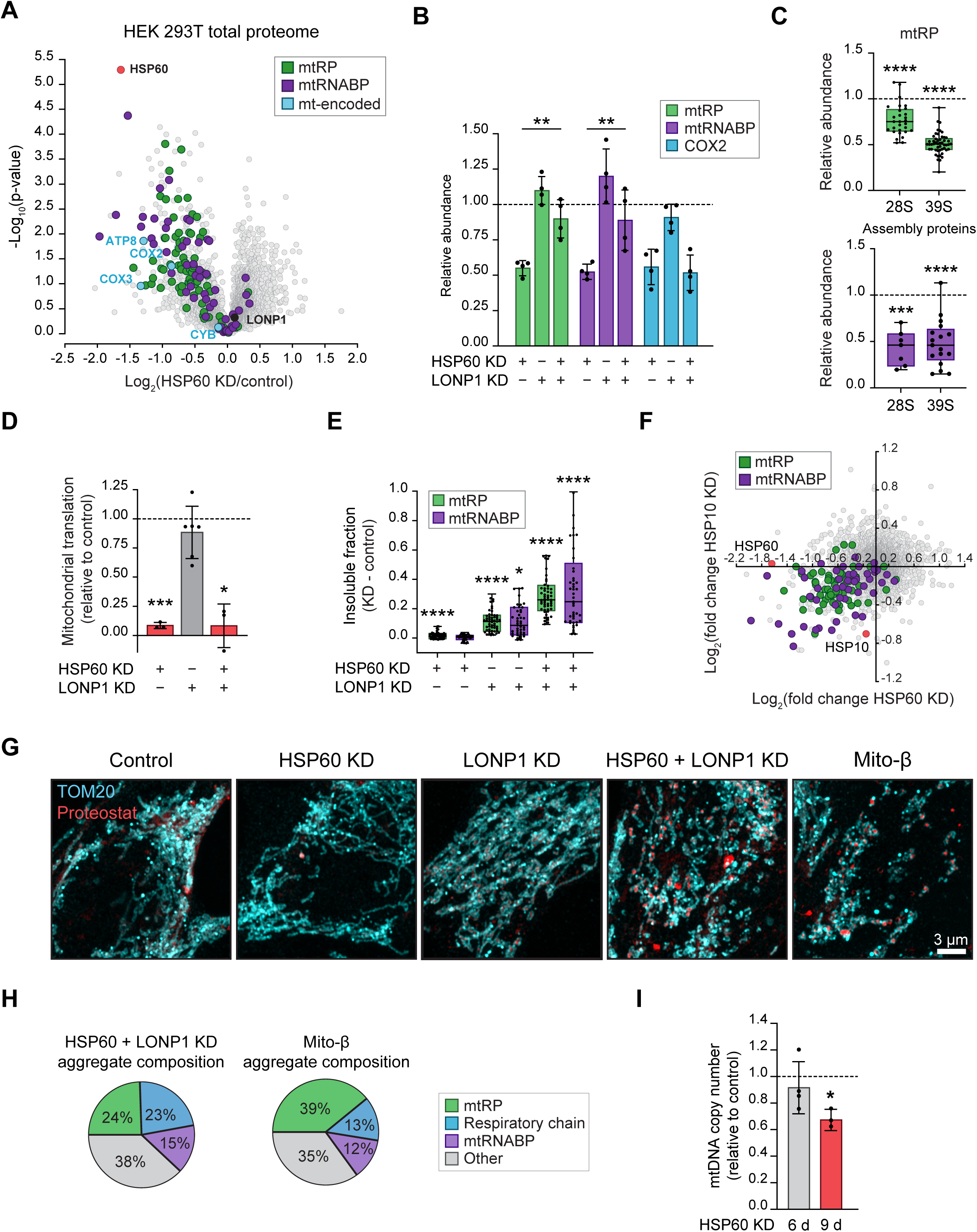
Disruption of mitochondrial proteostasis machinery phenocopies effects of protein aggregates. (A) Volcano plot of total proteome analysis of HEK 293T cells with HSP60 KD by label-free LC-MS/MS. mtRP, mtRNABP and mt-encoded proteins are highlighted. See also Table S4. (B) Summed abundance of the 20 most reduced mtRPs and mtRNABPs in HSP60 KD cells in (A) and of COX2 relative to LONP1 KD cells and to cells with combined HSP60 and LONP1 KD. Protein levels in WT cells are set to 1 (dotted line). Mean with error bars as SD (n = 4). p-Values by One-way ANOVA with Bonferroni′s multiple comparisons test. **p < 0.01. See also Table S4. (C) Mitoribosome assembly defect in HSP60 KD cells. Mitoribosomes in WT and HSP60 KD cells were fractionated by sucrose gradient centrifugation and analyzed by label-free LC-MS/MS as in Figure 3F (see also Figure S6B). Boxplot shows relative abundance of 28S and 39S mtRPs and ribosome assembly proteins in fractions corresponding to assembled mitoribosomes (summed LFQ values from fraction 4-7 for 28S and fraction 5-7 for 39S). Corresponding values in WT cells set to 1 (dotted line). Boxes and whiskers indicate quartiles and the line indicates the median (n = 3). p-Values by Mixed-effects analysis with Bonferroni′s multiple comparisons test. ***p < 0.001, ****p < 0.0001. See also Table S2. (D) Mitochondrial translation in HSP60 KD and LONP1 KD cells. Newly-synthesized mitochondrial proteins were labeled by HPG incorporation and Click chemistry as in Figure 4C. Quantification of mean fluorescence intensity from at least 15 cells per repeat is shown (see Figure S6C for representative images). Mean with error bars as SD (n ≥ 3). p-Values by Mixed-effects analysis with Dunnett′s multiple comparisons test. *p < 0.05, ***p < 0.001. (E) Analysis of insoluble mtRPs and mtRNABPs in HEK 293T cells upon HSP60 and LONP1 KD by label-free LC-MS/MS. Box plots show the insoluble fraction (pellet/total). Boxes indicate the 25th to 75th percentile, whiskers show the 10th to 90th percentile, and the line indicates the median. Background from control was subtracted (n = 4). p-Values by Mixed-effects analysis with Dunnett′s multiple comparisons test. *p < 0.05, ****p < 0.0001. See also Figures S6D, S6E and S6F and Table S4. (F) Comparative proteome analysis of HEK 293T cells with HSP10 KD (y-axis) and HSP60 KD (x-axis) by label-free LC-MS/MS. See also Table S4. (G) Visualization of protein aggregates in HEK 293T cells upon HSP60 and LONP1 KD or Mito-β expression by fluorescence microscopy. Mito-β expression was induced for 3 days with Dox. Aggregates were stained with Proteostat dye (red). Immunofluorescence staining of TOM20 with anti-TOM20 antibody was used to visualize mitochondria (cyan). (H) Aggregate composition by protein abundance in cells with combined LONP1 and HSP60 KD or Mito-β expression by LC-MS/MS. Aggregate composition was calculated based on the increase in abundance of mitochondrial proteins in the pellet fractions (iBAQ-values). The segmented pie chart shows the contribution of the major protein groups to total aggregate composition. See also Table S4. (I) Relative mitochondrial DNA (mtDNA) copy number in HEK 293T cells with HSP60 KD. Total DNA was isolated after 6 days and 9 days of KD, and quantified by qPCR. Mean with error bars as SD (n = 3). p-Values from Mixed-effects analysis with Bonferroni′s multiple comparisons test. *p < 0.05.

Surprisingly, analysis of the insoluble proteome of HSP60 KD cells revealed only a minor increase in insolubility of a limited number of mtRPs and mtRNABPs (Figures 6E and S6D; Table S4C). Given the ∼50% decrease in assembled mitoribosomes and breakdown of translation upon HSP60 KD (Figures 6C and 6D), we surmised that misfolded proteins might be degraded in HSP60 deficient cells. To test this hypothesis, we downregulated the mitochondrial matrix protease, LONP1, either alone or in combination with HSP60 (Figure S6A). LONP1 KD alone (by ∼50%) had no effect on mitochondrial translation (Figures 6D and S6C) but caused a limited but detectable increase in insoluble mtRP and RNABP proteins (Figures 6E and S6E). This is consistent with the observation of aggregates in mitochondria of cells from CODAS patients carrying a mutation in LONP1^40^ and confirms our finding (Figure 3E) that the efficiency of mitoribosome biogenesis is limited even in the absence of proteostatic stress. Strikingly, LONP1 KD substantially stabilized mtRPs and mtRNABPs in HSP60 KD cells almost to control levels (Figure 6B), resulting in extensive aggregation (Figures 6E and S6F). Note that COX2 levels did not increase, as mitochondrial translation remained impaired. Thus, LONP1 degrades dysfunctional mtRPs and mtRNABPs when HSP60 is deficient. Downregulation of the essential HSP60 cofactor, HSP10,^82^ also caused a reduction in the levels of most mtRPs and RNABPs (Figure 6F; Table S4D), suggesting that these proteins rely on transient encapsulation inside the HSP60 cavity by HSP10 for folding.^83^ The combined KD of HSP60 and LONP1, and to a lesser extent the KD of LONP1, resulted in the formation of intramitochondrial aggregate inclusions stainable with Proteostat dye (Figure 6G),^84^ phenocopying the inclusions formed by Mito-β (Figure 6G). Upon combined KD of HSP60 and LONP1, mtRPs, respiratory chain complex proteins and mtRNABPs contributed ∼24%, 23% and 15% to the insoluble fraction by abundance, respectively, similar to the composition of proteins co-aggregating with Mito-β (Figure 6H; Table S4E and S4F).

Interestingly, prolonged downregulation of HSP60 alone was sufficient to cause a decline in mtDNA copy number (Figure 6I) similar to the effect of the Mito-β aggregates (Figure 5D), consistent with mtRP deficiency (and consequent impairment of mitochondrial translation) being sufficient to drive mtDNA loss.

Taken together, these findings demonstrate the profound conformational vulnerability of the mitochondrial translation machinery under conditions that reflect impaired mitochondrial proteostasis in diseases caused by HSP60 or LONP1 mutations. Thus, defects in key components of mitochondrial protein quality control and mitochondrial protein aggregates have similar detrimental and mutually reinforcing consequences.

## DISCUSSION

In this study, we investigated the functional consequences of protein aggregation in yeast and human mitochondria, revealing an unexpected conformational vulnerability of the mitochondrial translation system to proteostatic stress. Import of Mito-β and OTCΔ into mitochondria generated large aggregate inclusions in the mitochondrial matrix. These inclusions locally disrupted inner membrane architecture and induced extensive co-aggregation and sequestration of newly-imported mtRPs and mtRNABPs (Figures 7A and 7B). As a result, mitoribosome assembly and mitochondrial translation were severely impaired, causing respiratory deficiency (Figure 7B) and the loss of mtDNA as a long-term consequence (Figure 7B). Consistent with the conformational susceptibility of the mitoribosome system, we found the efficiency of mitoribosome biogenesis to be limited even in the absence of proteostatic challenge, with multiple mtRPs and mtRNABPs requiring the chaperonin HSP60 for folding and assembly (Figure 7A). Failed mtRPs are normally degraded by the LONP1 protease, but undergo extensive aggregation when both HSP60 and LONP1 functions are reduced (Figure 7C). Thus, proteostatic stress in mitochondria, due to dysfunction of key quality control machinery or resulting from the formation of protein aggregates, disrupts mitochondrial protein synthesis and oxidative phosphorylation.

**Figure 7.**
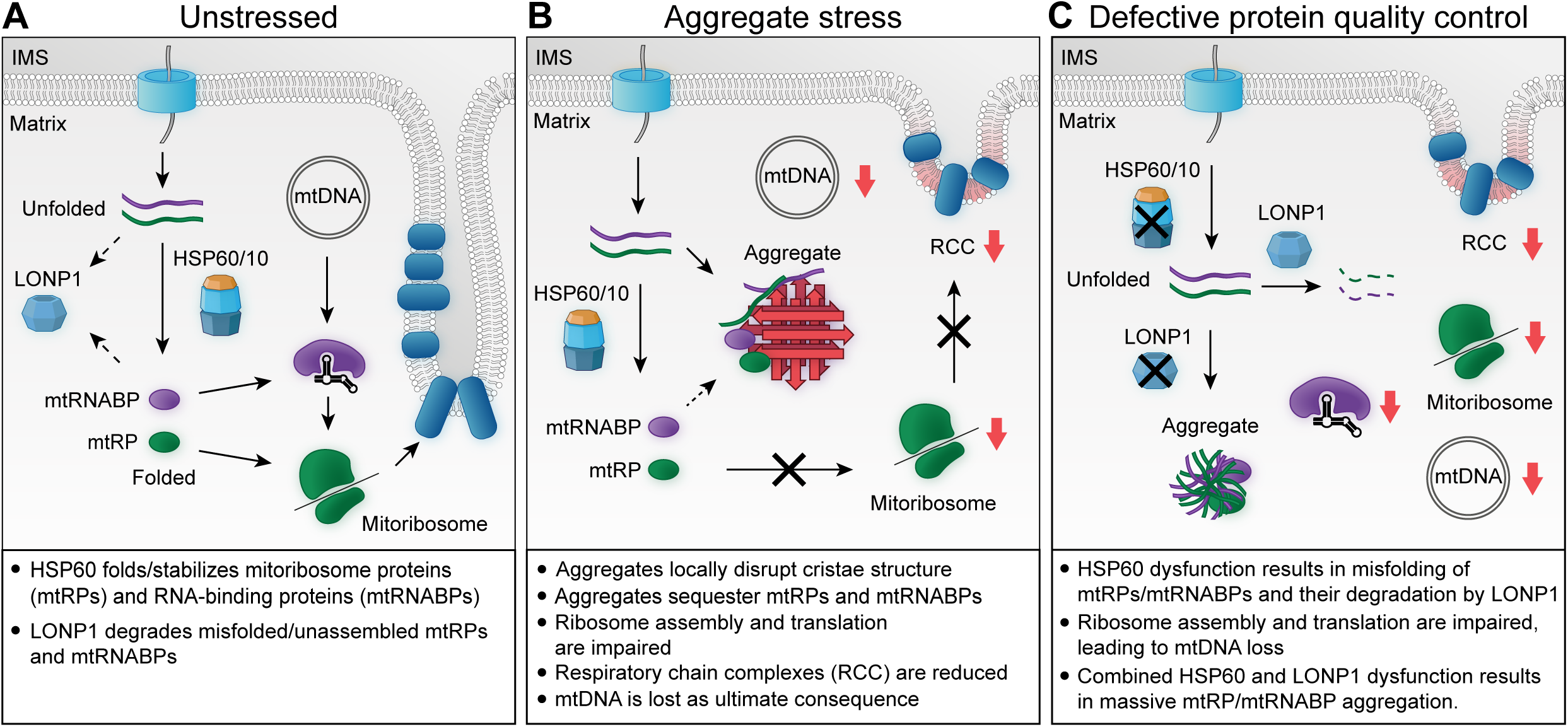
Vulnerability of the mitochondrial translation machinery to proteostatic stress. (A) Unstressed: A small set of respiratory chain proteins are encoded by mtDNA. These proteins are synthesized on mitochondrial ribosomes and inserted into the cristae of the inner mitochondrial membrane. All other mitochondrial proteins are imported as unfolded polypeptide chains and fold and assemble inside the organelle. mtRPs and mtRNABPs require the HSP60/HSP10 chaperonin system for folding/assembly. The efficiency of this reaction is limited, with misfolded or misassembled proteins being degraded by LONP1. (B) Aggregate stress: Protein aggregation inside the mitochondrial matrix generates large inclusions, locally disrupting cristae structure, and triggering a chain of detrimental events. The aggregates induce extensive co-aggregation and sequestration of newly-imported mtRPs and mtRNABPs. As a result, mitochondrial ribosome assembly and translation are impaired. This leads to a reduction in subunits of respiratory chain complexes (RCC) and prevents respiratory cell growth. Ultimately, mitochondrial DNA copy number decreases. (C) Defective protein quality control machinery: Upon HSP60 dysfunction, mtRPs and mtRNABPs are degraded by LONP1, but undergo aggregation when both HSP60 and LONP1 are reduced. As a result, mitochondrial protein synthesis and oxidative phosphorylation are severely impaired and mtDNA undergoes decline phenocopying the toxic effects of protein aggregates in (B).

### Mitoribosome biogenesis as target of aggregate toxicity

The toxic effects of protein aggregation vary substantially, depending on the specific cellular environment in which the aggregates form.^42–44,85,86^ Expression of the β-sheet model protein in the cytosol and nucleus has been shown to result in toxic aggregates, while in the ER the protein was maintained in a non-toxic gel-like state.^42–44,86^ In the present study, we observed that mitochondrial Mito-β and OTCΔ aggregate in the matrix, triggering a sequence of toxic events (Figure 7B). In addition to their large size, which apparently results in the local disappearance of cristae, the mitochondrial β-protein aggregates primarily exert toxic gain-of-function effects through aberrant interactions with mitochondrial proteins, leading to their co-aggregation and sequestration. While overall solubility was only moderately reduced for most proteins, co-aggregation was extensive and functionally relevant for mtRPs and other mtRNABPs (Figure 7B). As a consequence, fully assembled 28S ribosomal subunits were depleted by 40% and mitochondrial translation was reduced by 80-90% in human cells. It is noteworthy that this effect is specific for mitochondrial ribosome biogenesis, since aggregates of the same β-protein in the cytosol did not interact detectably with cytosolic ribosomal subunits.^42^

Why is mitoribosome biogenesis such a highly vulnerable process? All mtRPs (except Var1 in yeast) are nuclear encoded and must be imported from the cytosol into the mitochondrial matrix, where they fold and assemble with the mt-encoded ribosomal RNA in a pathway that relies on multiple specific assembly factors.^87–90^ Indeed, newly-imported ribosome proteins were preferentially targeted by the aggregates, presumably before completing folding and/or assembly. Importantly, several assembly factors, including the critical rRNA binding and processing proteins ERAL1 and MRPP3, were also sequestered. Thus, one explanation for the conformational sensitivity of mitoribosome biogenesis is that it occurs in the same compartment where most other imported proteins fold and assemble. In contrast, cytosolic ribosomes primarily assemble in the nucleus and nucleolus,^91^ thereby limiting the presence of unassembled ribosomal proteins in the cytosol. The high vulnerability of mitochondrial ribosome biogenesis may have increased the evolutionary pressure to transfer the vast majority of mitochondrial genes to the nucleus.

### Role of chaperonin HSP60 in mitoribosome biogenesis

Consistent with these considerations, we found the folding/assembly of mtRPs and other mtRNABPs to depend critically on the function of the chaperonin HSP60, with failed mtRPs being degraded by LONP1, but aggregating upon LONP1 dysfunction (Figures 7A and 7C). LONP1 has also been suggested to contribute to mtRP maturation in a chaperone-like manner, as strong downregulation (>90%) of LONP1 has been reported to cause mtRP aggregation.^92^ Our finding that the efficiency of mtRP folding/assembly is limited even under normal conditions, with LONP1 clearing misfolded mtRPs, provides an alternative explanation for this effect (Figures 3E and 7A). Other mitochondrial chaperones are apparently unable to replace the role of HSP60 in mitoribosome biogenesis, although reduced levels of mtRPs were also observed upon inhibition of mitochondrial HSP90 and HSP70.^29,93,94^ Thus, HSP60 (in conjunction with LONP1) could function as a nuclear-encoded regulator of the level of functional mitoribosomes and thus mitochondrial translation.

mtRPs and other mtRNABPs have previously been identified as HSP60 interactors in HEK 293T cells,^95^ attributed to their high abundance, but the critical role of HSP60 and its cofactor HSP10 in mitoribosome biogenesis had remained unknown. Ribosomal proteins and RNABPs are generally rich in unstructured sequence regions and, in their unassembled state, expose structurally dynamic interaction domains that may be highly susceptible to aberrant interactions.^96^ RNABPs also undergo binding-induced disorder-to-order transitions when they engage their substrates.^97^ These proteins may interact with the cylindrical HSP60 complex not only for initial folding, but also to be stabilized against aggregation prior to assembly. This may be achieved by transient encapsulation in the HSP60 cavity by HSP10 (Figure 7A). Notably, bacterial ribosomal proteins are not among the predominant clients of GroEL, the bacterial homolog of HSP60.^98,99^ The wall of the GroEL cavity in which protein folding occurs is negatively charged,^83^ which may result in incompatibility with the positively charged ribosomal proteins and RNABPs. Interestingly, this negative charge character is reduced in HSP60,^100^ perhaps representing an adaptation to this class of client proteins. We note that ribosomal subunits and assembly intermediates may generally require stabilization against aggregation, but for ribosomes of the eukaryotic cytosol this function is apparently provided by the specialized biocondensate environment of the nucleolus, consistent with the ability of the nucleolus to store and protect misfolded proteins during stress.^86^

Sequestration of newly-imported mtRPs and other mtRNABPs by Mito-β and OTCΔ aggregates may then occur in competition with binding and encapsulation by HSP60 (Figure 7B). Consistent with our findings, newly-imported proteins in mitochondria have been described to be particularly aggregation-prone under conditions of heat and ethanol stress.^101^ Although we identified HSP60 and several other proteostasis factors as aggregate interactors, the majority of HSP60 remained soluble, arguing against a direct interference of the aggregates with HSP60 function.

### mtDNA loss as ultimate consequence of proteostatic breakdown

As a late consequence of protein aggregation in mitochondria, we observed a decline of mtDNA content in yeast and human cells (Figure 7B). A gradual loss of mtDNA and functional capacity of the mitochondrial system is generally associated with aging.^3,5,10,11^ In yeast, loss of mtDNA occurs during replicative aging^67^ and may result from various defects in mitochondrial function, including deletion of mitochondrial ribosomal proteins, ATP-synthase subunits, mtDNA binding proteins, ISC biogenesis proteins, and LONP1.^11,73–76,92,102–104^ We identified multiple members of these protein groups, most prominently mtRPs, as aggregate interactors (Figure 7B). On the other hand, downregulation of HSP60 in HEK 293T cells, while disrupting mitoribosome biogenesis and mitochondrial translation, resulted in a clear reduction of mtDNA without causing overt protein aggregation (Figure 7C). Therefore, it is likely that the decline of mtDNA due to protein aggregation is caused indirectly by the loss of functional mtRPs. In yeast, the mtRP MHR1 and the ribosomal assembly protein MAM33 also function directly in mtDNA maintenance and mtDNA copy number regulation, respectively.^105,106^ Furthermore, changes in mtDNA copy number in humans have been associated with gene variants including LONP1, mtRPs and mtDNA binding proteins,^11^ supporting a link between mitochondrial proteostasis, mtRPs and mtDNA maintenance.

### Implications for diseases associated with mitochondrial proteostasis dysfunction

Loss-of-function mutations in HSP60 are associated with neurodegenerative conditions such as spastic paraplegia 13 (SPG13) and mitCHAP-60 disease.^38,39,107–110^ Our experiments mimicking HSP60 dysfunction by HSP60 knockdown suggest that defective mitochondrial translation, leading to respiratory deficiency, contributes to the cytopathology in these diseases (Figure 7C).

We found that mtRPs and mtRNABPs are degraded by LONP1 when HSP60 is defective (Figure 7C). Thus, LONP1 plays a major role in removing misfolded proteins that accumulate upon HSP60 dysfunction, reminiscent of the function of Lon protease in the bacterial cytosol.^111^ Importantly, LONP1 functions in the degradation of misfolded/misassembled mtRPs even under normal conditions (Figure 7A), explaining why protein aggregation is associated with LONP1 mutations that cause the developmental disease CODAS syndrome.^40^

Collapse of mitochondrial proteostasis, modeled by combined downregulation of HSP60 and LONP1, led to extensive aggregation of mtRPs and mtRNABPs (Figure 7C), the groups of proteins that are most susceptible to sequestration by pre-existing aggregates. Thus, the gain-of-function toxicity of protein aggregates phenocopies the consequences of proteostasis machinery breakdown. A decline in proteostasis capacity associated with aging^2,3^ would therefore be predicted to impair mitoribosome function. Indeed, aged worms contain fewer mitoribosomes and respiratory complex proteins^112^ and the brain of aged killifish shows reduced levels of basic proteins including mtRPs and RNABPs.^113^

In conclusion, the mitochondrial translation machinery is highly conformationally vulnerable. Therefore, respiratory dysfunction caused by degradation or aggregation of mtRPs and mtRNABPs is likely to contribute to the pathology of SPG13, MitCHAP-60, CODAS syndrome, and possibly other age-related diseases in which mitochondrial proteostasis is disrupted.

### Limitations of the study

Our findings identify the assembly of mitoribosomes as a process highly vulnerable to impairment of mitochondrial proteostasis, caused by targeting aggregation-prone proteins to the mitochondrial matrix or by downregulating the HSP60/LONP1 axis. However, the extent to which these observations are consistent with the effects of HSP60 and LONP1 mutations in patients remains to be determined. Furthermore, while our analysis of aggregate toxicity in mitochondria point to a clear link between the impairment of mitochondrial ribosomes and a decline in mtDNA, the exact sequence of steps leading to mtDNA loss and the underlying mechanism remains to be elucidated. Another aspect of future research is to determine how exactly mtRPs and mtRNABPs utilize the HSP60/HSP10 chaperonin during ribosome biogenesis. Do these proteins undergo folding while transiently being encapsulated in the chaperonin chamber, as demonstrated for other substrate proteins, or are they merely stabilized against aggregation prior to assembly? In vitro reconstitution experiments will be required to provide critical mechanistic insights.

## ACKNOWLEDGEMENTS

We thank Markus Oster, Giovanni Cardone and Martin Spitaler from the MPIB Imaging Facility (RRID:SCR_025739) for assistance with confocal and SIM microscopy, image analysis and flow cytometry, and Albert Ries for assistance with mass spectrometry. We acknowledge Gopal Jayaraj for insightful discussions and Chandhuru Jagadeesan for help with yeast methods. Cole Sitron is acknowledged for critically reading the manuscript. We thank the Martin Ott lab for sharing the yeast reporter strain for mitochondrial translation. Mutant ornithine transcarbamylase (OTCΔ) was a gift from Nicholas Hoogenraad. This work was supported by funding from the Max-Planck-Förderstiftung, by the Deutsche Forschungsgemeinschaft (DFG, German Research Foundation) – SPP2453 project number 541592768, and by DFG under Germany’s Excellence Strategy within the framework of the Munich Cluster for Systems Neurology (EXC 2145 SyNergy – ID 390857198) and the “Multiscale Bioimaging: from Molecular Machines to Networks of Excitable Cells” (MBExC) Excellence Cluster (EXC 2067/1-390729940).

## AUTHOR CONTRIBUTIONS

H.H. designed and performed most experiments and supervised C.F.G. and A.D.. V.A.T. and K.E. performed cryo-electron tomography, supervised by R.F.-B.. C.F.G. performed toxicity assays, COX2 immunoblotting, ISR quantification and mtDNA microscopy in human cells. I.S. produced viruses and generated stable cell lines. I.P. and O.K.W. performed DNA-PAINT super-resolution microscopy experiments, supervised by R.J.. A.D. performed OTCΔ-interactome analysis. B.H. performed RNA-sequencing, supervised by R.S.. P.Y. performed differentiation of iPS-cells to forebrain-type neurons. R.K. operated the mass spectrometer. M.S.H. and F.U.H. initiated and supervised the project. F.U.H and H.H. wrote the manuscript with input from the other authors.

---

**Figure S1.**
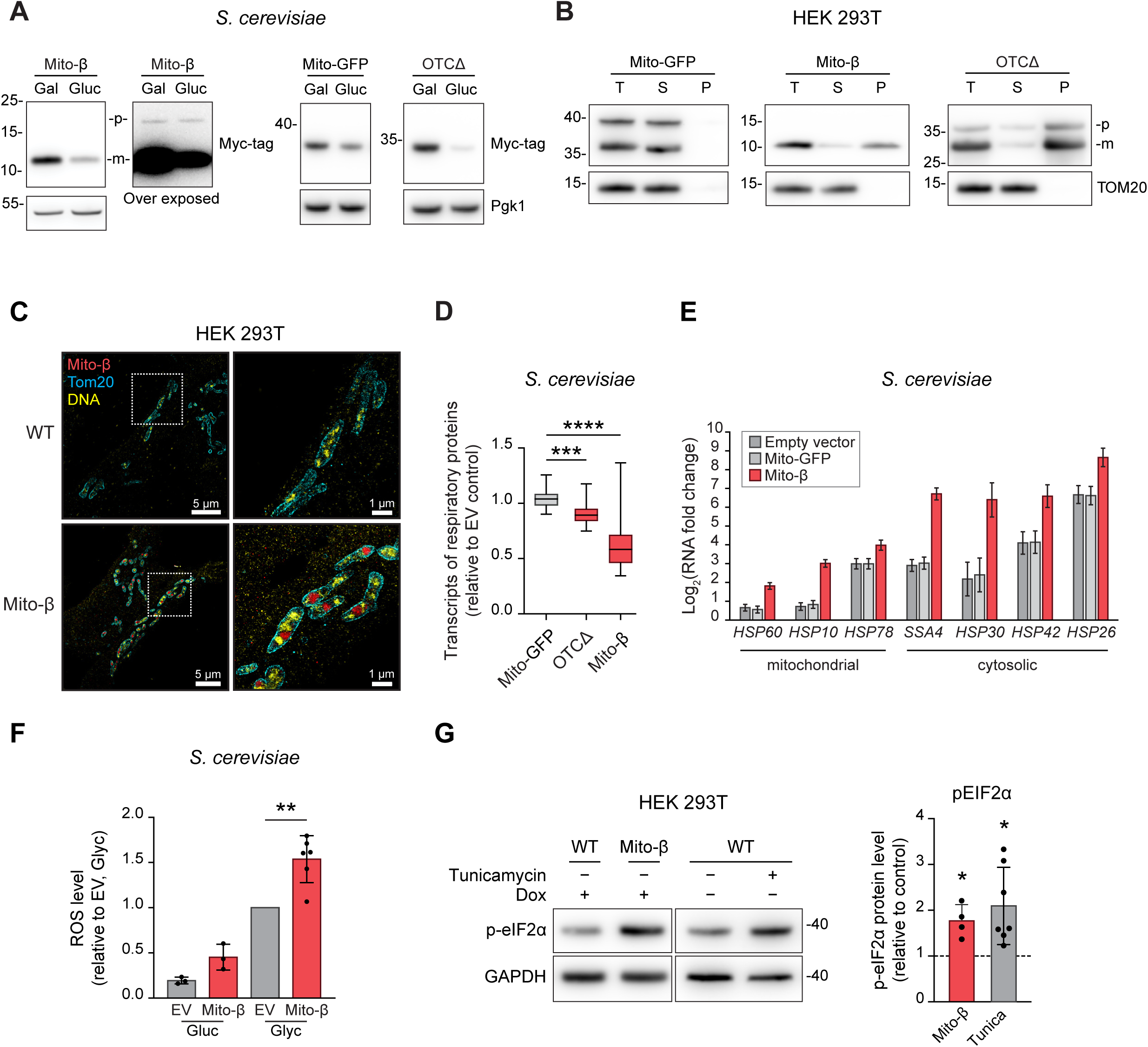
Mitochondrial protein aggregates impair respiratory function, related to Figure 1. (A) Levels of Mito-β, Mito-GFP and OTCΔ in yeast cells upon induction with galactose (Gal) or constitutive expression in glucose (Gluc) medium. Mito-β, Mito-GFP and OTCΔ were detected by immunoblotting with anti-myc antibody. Over-exposed image shows the Mito-β precursor band (p) in addition to the mature band (m). 3-Phosphoglycerate kinase (PGK1) served as loading control. (B) Solubility of Mito-GFP, Mito-β and OTCΔ in HEK 293T cells. 24 h after transfection, cells were lysed in RIPA buffer and lysates fractionated by centrifugation, followed by immunoblotting with anti-myc antibody. Outer mitochondrial membrane protein TOM20, detected with anti-TOM20 antibody, served as mitochondrial loading control. T: total, S: soluble, P: pellet, p: precursor, m: mature. (C) Immunofluorescence staining of mtDNA and Mito-β in HEK 293T cells by DNA-PAINT super resolution microscopy. WT cells and cells expressing Dox-inducible Mito-β were cultured with Dox for 24 h. Mito-β was detected with anti-myc antibody (red), mtDNA with anti-DNA antibody (DNA) (yellow), and TOM20 with anti-TOM20 antibody (cyan). (D) Analysis of RNA upregulation of respiratory chain complex subunits by RNAseq in yeast cells. Cells constitutively expressing Mito-GFP, Mito-β or OTCΔ were compared to EV control. RNA was isolated 5 h after shifting the cells from glucose to glycerol medium. Boxes and whiskers indicate quartiles and the line indicates the median (n = 3). p-Values from One-way ANOVA with Dunnett’s multiple comparison test. ***p < 0.001, ****p < 0.0001. (E) Change in RNA levels of selected cytosolic and mitochondrial chaperone genes as analyzed in (D). (F) Quantification of reactive oxygen species (ROS). Yeast cells constitutively expressing Mito-β and EV control cells grown in glucose (Gluc) or glycerol (Glyc) medium were incubated with 5 µM CellROX for 1 h and fluorescence intensity was quantified by flow cytometry. Values were normalized to EV control in glycerol medium. Mean with error bars as SD (n ≥ 3). p-Values by Mixed-effects analysis with Bonferroni′s multiple comparisons test. **p < 0.01. (G) Analysis of integrated stress response (ISR) protein p-eIF2α in HEK 293T cells. Mito-β expression was induced with Dox for 3 days. Cells were treated with 5 µg/mL tunicamycin (Tunica) for 5 h as positive control. Glycerinaldehyd-3-phosphat-dehydrogenase (GAPDH) served as loading control. Cell lysates were analyzed by immunoblotting with anti-p-eIF2α antibody (left panel), followed by quantification by densitometry (right panel). Levels of p-eIF2α of Mito-β-expressing cells and tunicamycin treated cells are shown relative to Dox treated control cells and untreated cells, respectively. Mean with error bars as SD (n ≥ 3). p-Values by Mixed-effects analysis with Bonferroni′s multiple comparisons test. *p < 0.05.

**Figure S2.**
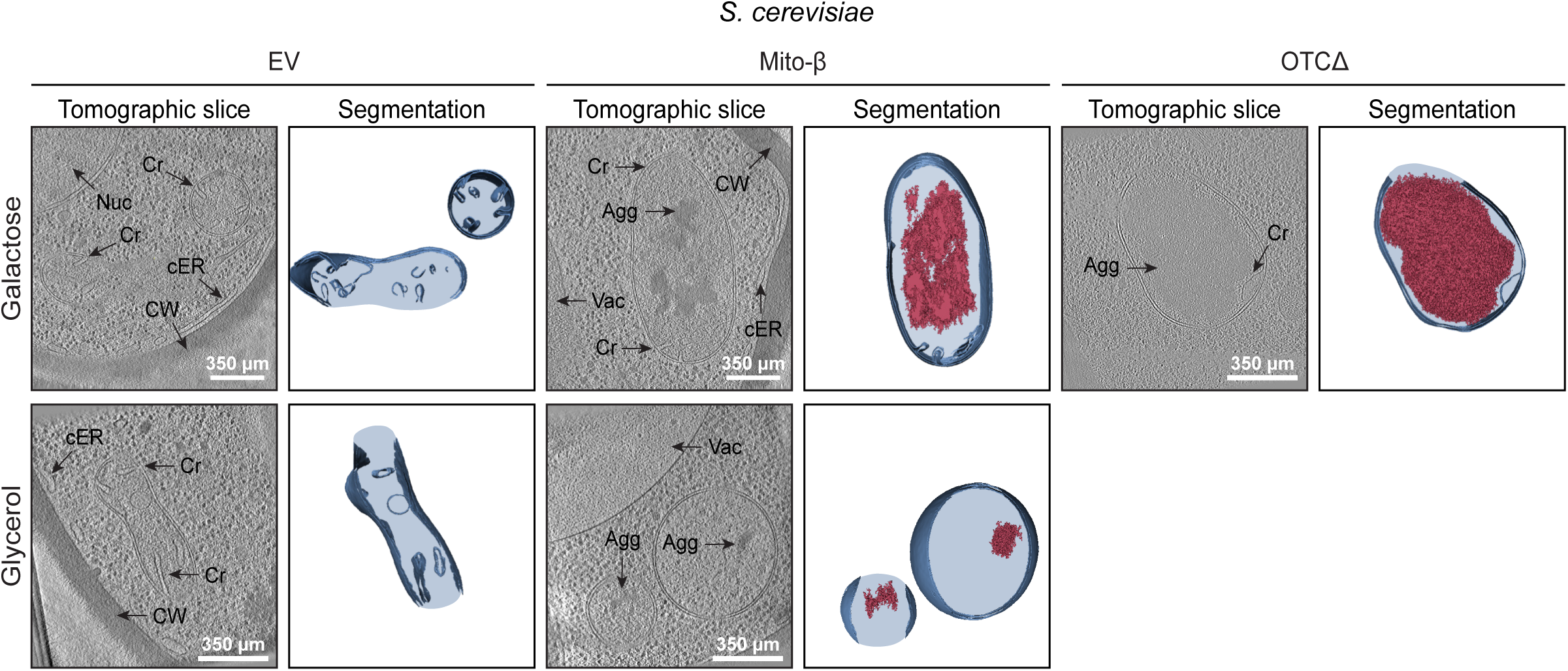
Protein aggregates impair mitochondrial cristae architecture, related to Figure 2. Cryo-electron tomographic slices (1.4 to 1.8 nm thickness) of yeast cells expressing galactose-inducible Mito-β and OTCΔ in galactose medium and constitutively expressing Mito-β in glycerol medium in comparison to EV control cells. Segmentations show mitochondrial membranes including cristae (Cr) in blue and aggregates (Agg) in red. Other visible cellular structures include nucleus (Nuc), cell wall (CW), vacuole (Vac) and cortical ER (cER). Tomograms are representative of at least 3 individual cells from 2 biological replicates. Scale bar: 350 µm

**Figure S3.**
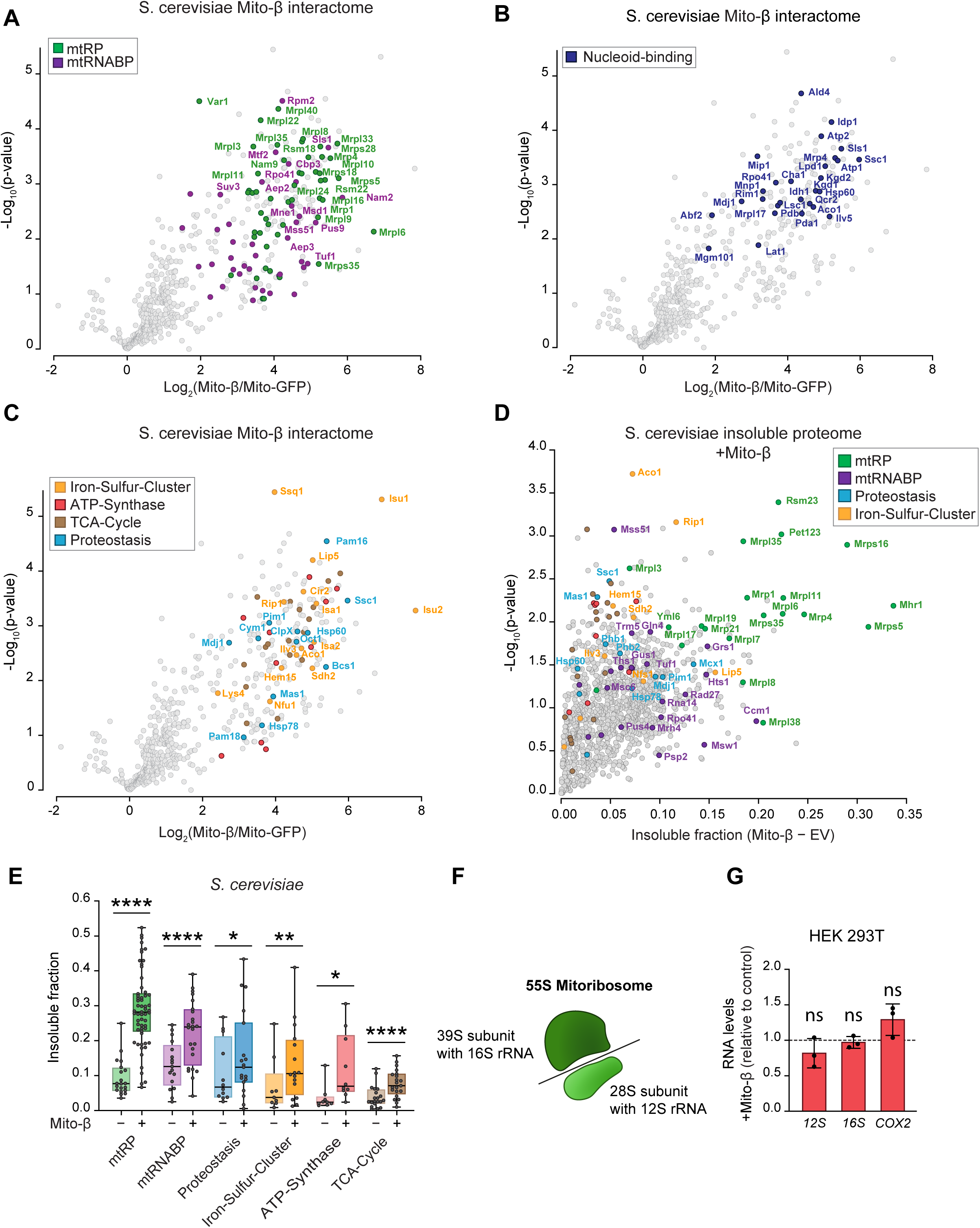
Mitochondrial aggregates sequester mtRPs, related to Figure 3. (A-C) Interactome analysis of Mito-β in yeast cells by SILAC LC-MS/MS, related to Figure 3A. Volcano plot of proteins co-immunoprecipitating with Mito-β in comparison to Mito-GFP after Mito-β/Mito-GFP expression for 24 h in galactose medium (from SILAC-ratios). Selected groups of proteins are highlighted: (A) mtRPs and mtRNABPs, (B) nucleoid-binding proteins, (C) proteostasis factors, iron-sulfur-cluster proteins, ATP-synthase subunits and TCA cycle proteins. Average fold changes (n = 3). p-Values by Students t-test. See also Table S1. (D) Analysis of the insoluble proteome of Mito-β-expressing yeast cells by SILAC LC-MS/MS. Average insoluble fraction (pellet/total, from SILAC ratios) of the proteins indicated is plotted against p-value by Students t-test. Background from EV control was subtracted. Selected groups of proteins are color coded. See also Table S1. (E) Analysis of the insoluble proteome of Mito-β-expressing yeast cells. Insoluble fraction (pellet/total, from SILAC-ratios) of protein groups in (D) is plotted. Boxes and whiskers indicate quartiles and the line indicates the median (n=3). p-Values by Mixed-effects analysis with Bonferroni′s multiple comparisons test. *p < 0.05, **p < 0.01, ****p < 0.0001. See also Table S1. (F) Schematic representation of the 55S mitoribosome consisting of the large 39S subunit with the 16S rRNA and the small 28S subunit with the 12S rRNA. (G) Quantification of *12S*, *16S* and *COX2* RNA in WT and HEK 293T cells expressing Mito-β for 3 days with Dox by qPCR. Mean with error bars as SD (n = 3). No significant results by Repeated-measures One-way ANOVA with Dunnett′s multiple comparisons test.

**Figure S4.**
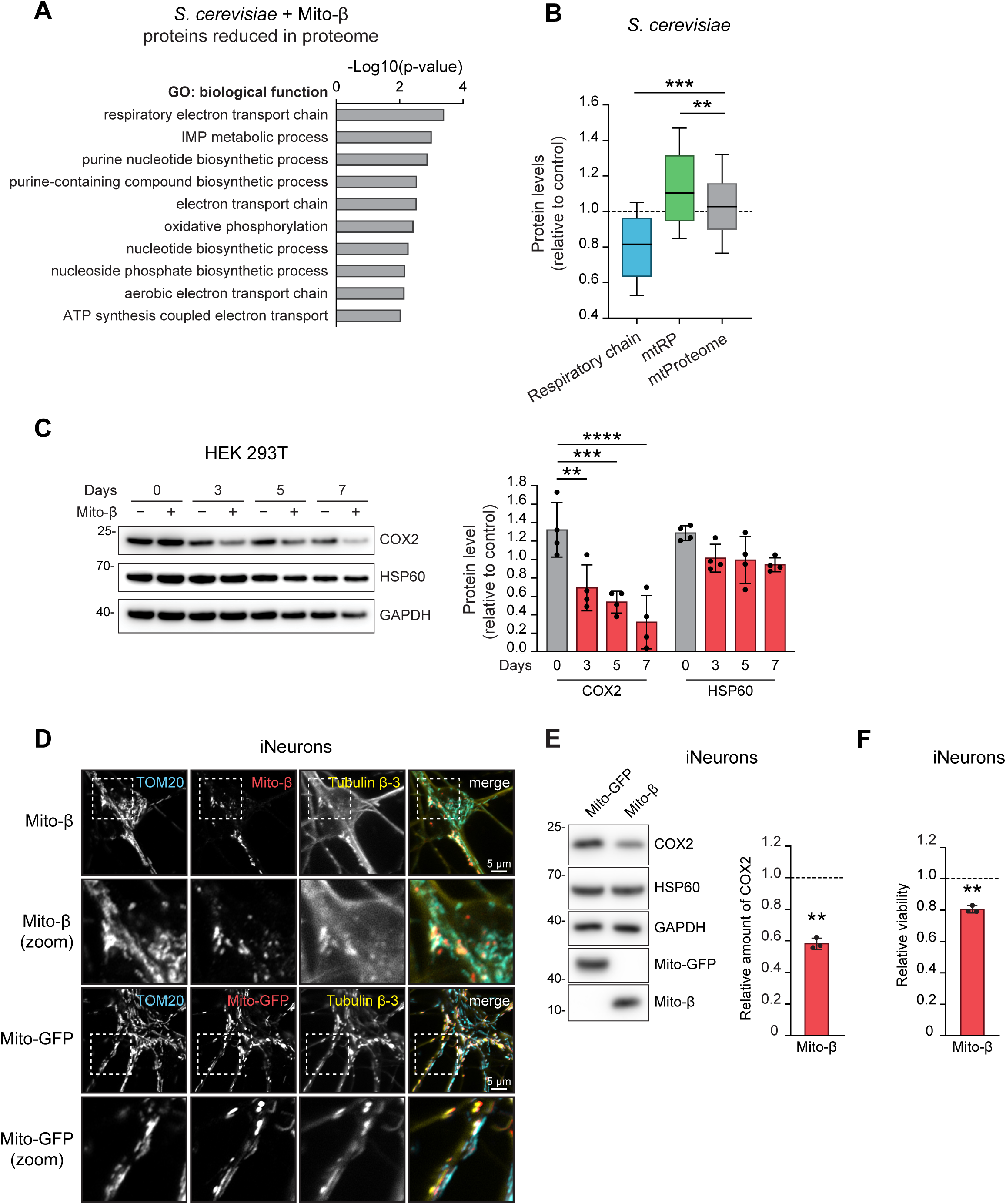
Mitochondrial aggregates impair mitochondrial translation, related to Figure 4. (A) Proteome analysis of Mito-β-expressing yeast cells displayed as GO-term enrichment of proteins that are at least 25% reduced in abundance compared to WT cells (see also Figure 4B and Table S3). (B) Proteome analysis of Mito-β-expressing yeast cells. Relative protein amount of respiratory chain proteins, mtRPs and all mitochondrial proteins (see also Figure 4B and Table S3). Boxes indicate the 25th to 75th percentile, whiskers show the 10th to 90th percentile, and the line indicates the median (n = 4). p-Values by One-way ANOVA with Dunnett’s multiple comparison test. **p < 0.01, ***p < 0.001. (C) Analysis of COX2 level in HEK 293T cells with Mito-β expression. WT cells and cells expressing Dox-inducible Mito-β were cultured for up to 7 days in the presence of Dox, followed by anti-COX2 immunoblotting (left panel). GAPDH and HSP60 were analyzed as loading control. Quantification of protein amounts by densitometry is shown as bar graph in Mito-β-expressing cells relative to control (right panel). Mean with error bars as SD (n = 4). p-Values by One-way ANOVA with Bonferroni′s multiple comparisons test. **p < 0.01, ***p < 0.001, ****p < 0.0001. (D) Immunofluorescence staining of iNeurons expressing Mito-β and Mito-GFP. iNeurons were cultured for 7 days after lentiviral transduction. Mito-β and Mito-GFP were detected with anti-myc antibody (myc, red); anti-TOM20 antibody staining visualizes mitochondria (cyan) and anti-Tubulin β-3 staining was used as neuronal marker (yellow). (E) Analysis of COX2 levels in iNeurons with Mito-β expression. iNeurons were cultured for 14 days after lentiviral transduction. Mito-β and Mito-GFP were detected by immunoblotting with anti-myc antibody. GAPDH and HSP60 served as loading control (left panel). Quantification of protein amounts by densitometry in Mito-β-expressing cells relative to Mito-GFP expressing cells is shown as bar graph (right panel). Mean with error bars as SD (n = 3). p-Values by One-sample Student’s t-test. **p < 0.01. (F) Analysis of cellular viability by MTT-assay in iNeurons with Mito-β expression. iNeurons were cultured for 14 days after lentiviral transduction. Viability of Mito-β-expressing cells is shown relative to Mito-GFP expressing cells. Mean with error bars as SD (n = 3). p-Values by One-sample Student’s t-test. **p < 0.01.

**Figure S5.**
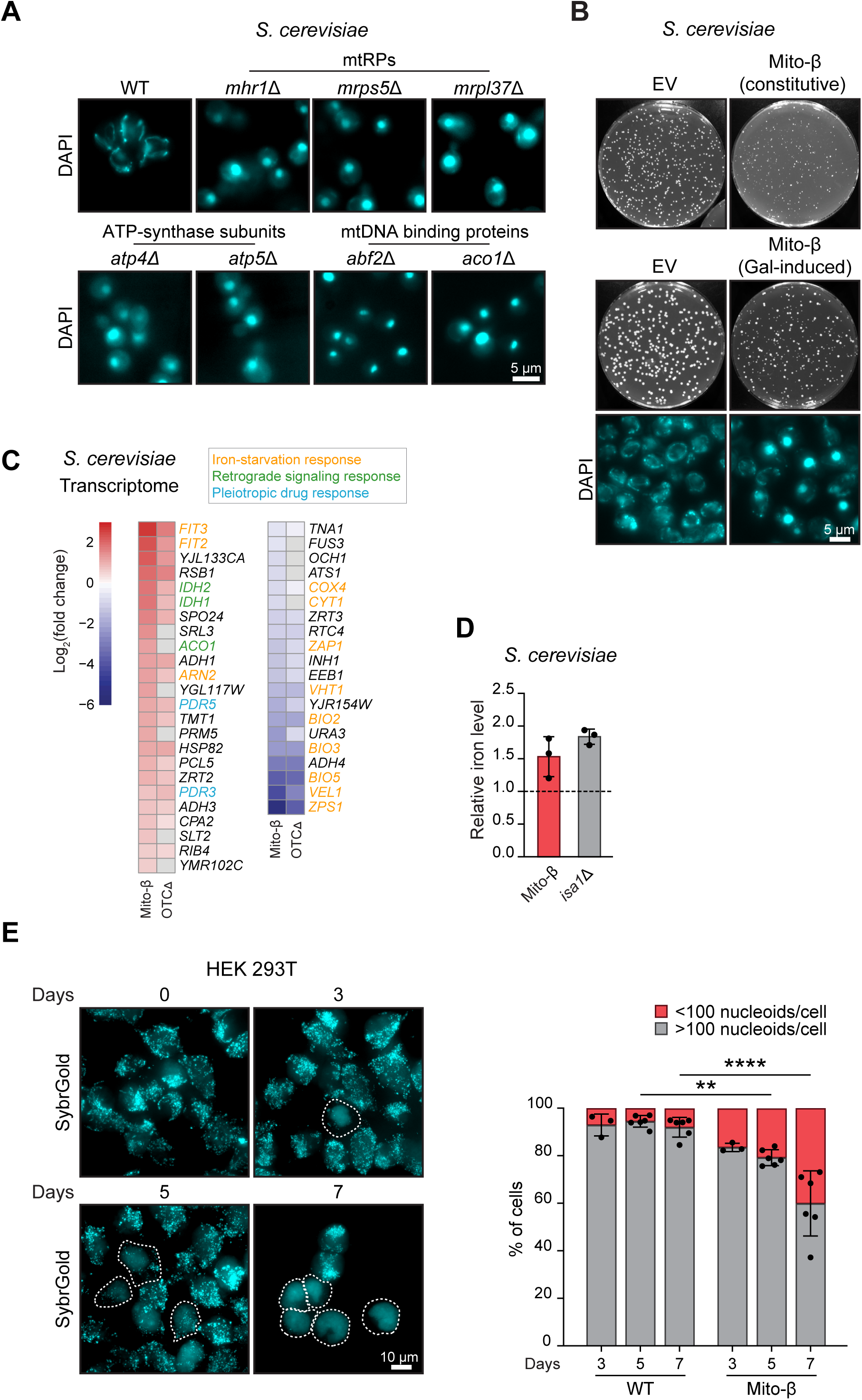
Mitochondrial aggregates cause mtDNA loss, related to Figure 5. (A) Visualization of mtDNA loss in yeast. WT cells and cells deleted in specific mtRPs (*mhr1*Δ, *mrps5*Δ, *mrpl37*Δ), ATP-synthase subunits (*atp4*Δ, *atp5*Δ) and mtDNA binding proteins (*abf2*Δ, *aco1*Δ) were grown for 24 h in glucose medium, stained with DAPI and analyzed by live cell fluorescence microscopy. (B) Evaluation of colony size by plating of yeast cells with galactose-induced Mito-β expression. EV control and galactose-induced Mito-β-expressing cells were first grown on galactose plates for 2 to 3 days. Thereafter, 1/10^3^ of one picked colony was re-plated on galactose plates. Small colony size indicates petite phenotype. mtDNA was stained with DAPI in cells from the second plating. (C) Transcriptome analysis of yeast cells with mitochondrial aggregates. RNA was isolated from EV control cells and cells constitutively expressing Mito-β or OTCΔ grown in glucose medium for 24 h. Heat map displays significantly changed transcripts in Mito-β-expressing cells and corresponding values from OTCΔ expressing cells. Transcripts corresponding to the iron-starvation response (orange), retrograde signaling response (green) and pleiotropic drug response (cyan) are highlighted. (D) Quantification of cellular iron by colorimetric bathophenanthrolinedisulfonic acid assay. Yeast cells with constitutive Mito-β expression, EV control cells and ISA1 deletion cells (*isa1*Δ) as positive control were grown in glucose medium for 24 h. Cellular iron was quantified and displayed as bar graph relative to EV control. Mean with error bars as SD (n = 3). Iron content of WT cells is set to 1. (E) Visualization of mtDNA in HEK 293T cells. Mito-β expression was induced for up to 7 days with Dox. mtDNA was stained with SybrGold and visualized by live cell fluorescence microscopy. Boundaries of cells with partial and severe loss of mtDNA are outlined (see also Figure 5C). Cells (≥377 cells per condition) were classified in groups with <100 or >100 nucleoids per cell. The relative distribution of cells between these groups is displayed as a stacked bar graph. Mean with error bars as SD (n ≥ 3). p-Values by One-way ANOVA with Bonferroni′s multiple comparisons test. **p < 0.01, ****p < 0.0001.

**Figure S6.**
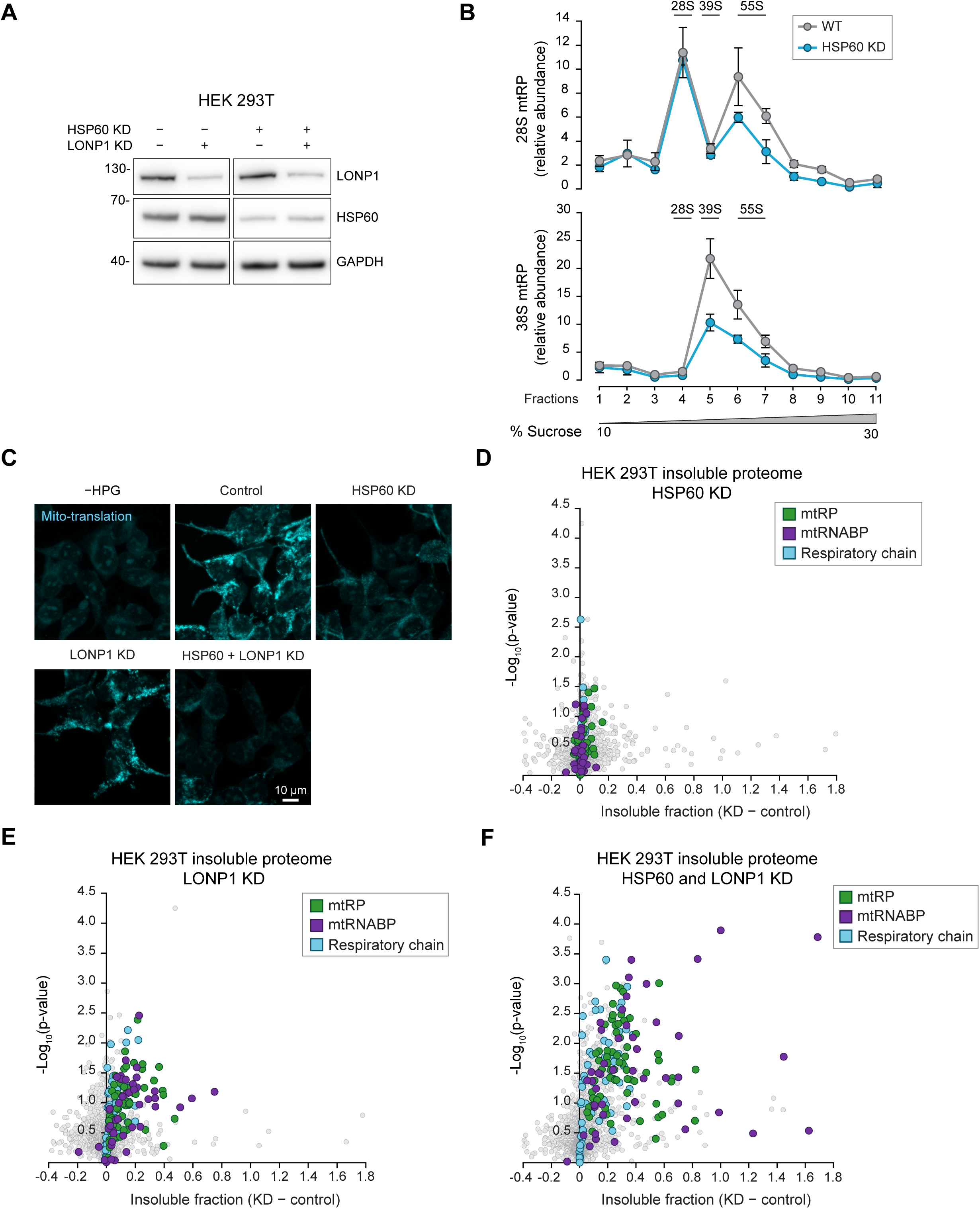
Disruption of mitochondrial proteostasis machinery phenocopies effects of protein aggregates, related to Figure 6. (A) Verification of HSP60 and LONP1 knockdown (KD) by Immunoblotting. HEK 293T cells expressed IPTG-inducible shRNA against HSP60 and scrambled shRNA as control for 6 days. Additionally, cells were transfected with siRNA against LONP1 and non-targeted (Nt) siRNA as control for the last 3 days. GAPDH served as loading control. (B) Analysis of mitoribosome assembly by sucrose gradient fractionation of HSP60 KD cells, followed by label-free LC-MS/MS of gradient fractions. Average abundance (summed LFQ-values) of 28S and 39S mtRPs per fraction is plotted (n=3) see also Table S2. (C) Mitochondrial translation in HSP60 KD and LONP1 KD cells. Fluorescence images show newly-synthesized mitochondrial proteins that were labeled by HPG incorporation and Click chemistry as in Figure 4C. (D-F) Analysis of the insoluble proteome of HEK 293T cells with HSP60 KD (C), LONP1 KD (D) and combined HSP60 and LONP1 KD (E) by label-free LC-MS/MS. The average insoluble fraction (pellet/total, from LFQ-values) (n=4) of individual proteins is plotted against p-value by Students t-test. Background from control cells was subtracted. Selected groups of proteins are highlighted. see also Table S4.

## SUPPLEMENTAL EXCEL TABLES

**Table S1. Interactome of Mito-β and OTCΔ aggregates and protein solubility in Mito-β-expressing cells, related to Figures 3A-E and S3A-E.**

(A) Interactome analysis of Mito-β in yeast cells by SILAC LC-MS/MS. Analysis of proteins co-immunoprecipitating with Mito-β in comparison to Mito-GFP and empty vector (EV) control after Mito-β/Mito-GFP expression for 24 h in galactose medium. Detected proteins are sorted according to their log_2_ fold change (Mito-β vs. Mito-GFP) derived from the log_2_ fold change (Mito-β vs. Empty vector) and log_2_ fold change (Mito-GFP vs. Empty vector), as based on their SILAC ratios across 3 independent biological replicates. p-Value by Students t-test. Related to Figures 3A, 3B and S3A-C.

(B) Interactome analysis of OTCΔ in yeast by SILAC LC-MS/MS. Analysis of proteins co-immunoprecipitating with OTCΔ in comparison to EV control after OTCΔ expression for 24 h in galactose medium. Detected proteins are sorted according to their log_2_ fold change (OTCΔ vs. EV) derived from their SILAC ratios across 3 independent biological replicates. p-Value by Students t-test. Related to Figure 3B.

(C) Analysis of the insoluble proteome of Mito-β-expressing yeast cells by SILAC LC-MS/MS. Cells were grown for 24 h in galactose medium to induce expression of Mito-β. Cell lysates were separated into total and insoluble fractions by centrifugation. Detected proteins are sorted according to their insoluble fraction (Mito-β subtracted by EV) derived from SILAC ratios, which represent pellet/total ratios across 3 independent biological replicates. p-Value by Students t-test. Related to Figures S3D and S3E.

(D) Analysis of the insoluble proteome of Mito-β-expressing HEK 293T cells by label-free LC-MS/MS. Wildtype (WT) cells and cells containing Dox-inducible Mito-β were cultured for 3 days with Dox. Cell lysates were separated into total and insoluble fraction by centrifugation. Detected proteins are sorted according to their insoluble fraction (Mito-β subtracted by wildtype (WT)) derived from LFQ values of pellet and total fractions, across 4 independent biological replicates. p-Value by Students t-test. Related to Figures 3C and 3D.

(E) Analysis of solubility of newly-imported proteins in Mito-β-expressing HEK 293T cells by SILAC LC-MS/MS. Mito-β expression was induced for 16 h in HEK 293T before cells were grown in heavy amino acid isotope for 4 h to label newly-translated proteins. Cell lysates were separated into total and insoluble fraction by centrifugation. The table displays the insoluble fraction of isotope-labeled and non-labeled proteins in WT and Mito-β-expressing cells. Insoluble fractions are derived from LFQ values of pellet and total fractions across 5 independent biological replicates. See also STAR Methods. Related to Figure 3E.

**Table S2. Analysis of mitoribosome assembly, related to Figures 3F, 3G, 6C and S6B.**

(A) Analysis of mitoribosomes in sucrose gradient fractions of Mito-β-expressing HEK 293T cells. WT cells and cells containing Dox-inducible Mito-β were cultured for 3 days with Dox. Mitochondrial lysates were fractionated into 11 fractions and analyzed by label-free LC-MS/MS. The table displays the protein abundance of the sucrose gradient fractions, the preclearance pellet fractions and total fractions of WT, Mito-β-expressing and HSP60 KD cells, derived from LFQ values across 3 independent biological replicates. See also STAR Methods. Related to Figures 3F, 3G, 6C and S6B.

**Table S3. Proteome analysis of Mito-β-expressing yeast cells, related to Figures 4B and S4A and S4B.**

(A) Proteome analysis of Mito-β-expressing yeast cells by label-free LC-MS/MS. Cells were grown for 24 h in galactose medium to induce expression of Mito-β. Detected proteins are sorted according to their log_2_ fold change (Mito-β vs. EV) derived from LFQ values across 4 independent biological replicates. p-Value by Students t-test. Related to Figures 4B, S4A and S4B.

**Table S4. Proteome analysis of HSP60, LONP1 and HSP10 KD cells and protein solubility in HSP60 and LONP1 KD cells, related to Figures 6A, 6B, 6E, 6F, 6H and S6D-F.**

(A) Proteome analysis of HEK 293T cells with HSP60 KD by label-free LC-MS/MS. HSP60 was downregulated by IPTG-inducible shRNA for 6 days. Detected proteins are sorted according to their log_2_ fold change (HSP60 KD vs. scrambled control) derived from LFQ values across 4 independent biological replicates. p-Value by Students t-test. Related to Figure 6A.

(B) Proteome analysis of HEK 293T cells with HSP60 and LONP1 KD by label-free LC-MS/MS. HSP60 was downregulated by IPTG-inducible shRNA for 6 days and LONP1 by siRNA for 3 days. The table shows protein fold changes of cells with HSP60 KD, LONP1 KD and combined HSP60 and LONP1 KD compared to control cells expressing scrambled shRNA or non-targeted siRNA, derived from LFQ values across 4 independent biological replicates. Related to Figure 6B.

(C) Analysis of the insoluble proteome of HEK 293T cells with HSP60 and LONP1 knockdown by label-free LC-MS/MS. HSP60 was downregulated by IPTG-inducible shRNA for 6 days and LONP1 by siRNA for 3 days. Cell lysates were separated into total and insoluble fractions by centrifugation. The table shows the insoluble fraction of proteins in cells with HSP60 KD, LONP1 KD and combined HSP60 and LONP1 KD compared to control cells expressing scrambled shRNA or non-targeted siRNA, derived from LFQ values of pellet and total fractions across 4 independent biological replicates. Related to Figures 6E and S6D-F.

(D) Proteome analysis of HEK 293T cells with HSP10 KD by label-free LC-MS/MS. HSP10 was downregulated by siRNA for 3 days. Detected proteins are sorted according to their log_2_ fold change (HSP10 KD vs. non-targeted control) derived from LFQ values across 3 independent biological replicates. p-Value by Students t-test. Related to Figure 6F.

(E-F) Analysis of aggregate composition of HEK 293T cells with HSP60 and LONP1 knockdown (E) or with Mito-β expression (F) by label-free LC-MS/MS. Protein abundance is based on iBAQ values across 4 independent biological replicates. Aggregate composition was calculated based on the increase in abundance of mitochondrial proteins in the pellet fractions. Related to Figure 6H.

## STAR METHODS

### Lead contact and resource availability

Further information and requests for resources and reagents should be directed to and will be fulfilled by the lead contact, F. Ulrich Hartl.

### Materials availability

Plasmids and strains generated in this study are available upon request to the lead contact.

### Data availability

The mass spectrometry proteomics data have been deposited to the ProteomeXchange Consortium^114^ via the PRIDE^115^ partner repository with the dataset identifier PXD059912. All transcriptome data generated in this study have been deposited in GEO and are available under accession number GSE295389. Raw data of immunoblots were deposited on Mendeley at (https://data.mendeley.com/preview/fbj6fhcd4c?a=e48ee97a-75cb-4bda-95c3-f78c040c671e). This paper does not report original code.

### Experimental model and subject details

#### Yeast culture

The S. cerevisiae strain used in this study was BY4741 (EUROSCARF). Genotypes of strains utilized, and genetically manipulated derivatives thereof, are listed in KEY RESOURCES TABLE. Cells were grown in synthetic dropout medium (6.7 g/L yeast nitrogen base without amino acids, 0.015 g/L adenine, 0.058 g/L alanine, 0.058 g/L arginine, 0.058 g/L asparagine, 0.058 g/L aspartic acid, 0.058 g/L cysteine, 0.058 g/L glutamine, 0.058 g/L glutamic acid, 0.058 g/L glycine, 0.058 g/L histidine, 0.058 g/L inositol, 0.058 g/L isoleucine, 0.292 g/L leucine, 0.058 g/L lysine, 0.058 g/L methionine, 0.006 g/L para-aminobenzoic acid, 0.058 g/L phenylalanine, 0.058 g/L proline, 0.058 g/L serine, 0.058 g/L threonine, 0.058 g/L tryptophan, 0.058 g/L tyrosine, 0.058 g/L uracil, 0.058 g/L valine) at 30 °C containing 2% glucose, unless otherwise indicated (glucose medium). For *GAL1-*induced over expression, cells were first grown in medium containing 2% raffinose (raffinose medium), before shifting to medium containing 3% galactose and 1% raffinose (galactose medium). For non-fermentative conditions, cells were grown in medium containing 3% glycerol (glycerol medium). Yeast cells were quantified in OD600 units, where 1 OD corresponds to the number of cells in 1 mL of culture with an OD600 of 1.

### Human cell culture

HEK 293T (ATCC, CRL-3216, RRID: CVCL_0063) and HeLa cells (ATCC, CCL-2, RRID: CVCL_0030) were cultured in DMEM (Gibco) supplemented with 10% FBS (Gibco), 1% penicillin/streptomycin (Gibco), 110 mg/L pyruvate (Gibco) and 4.5 g/L glucose (Gibco) (glucose medium), or 1.8 g/L galactose (Sigma-Aldrich) (galactose medium) at 37 °C in 5% CO_2_.

### iPSC-derived neurons (iNeurons)

The induced pluripotent stem cell (iPSC) line HPSI0214i-kucg_2 was purchased from the UK Health Security Agency (#77650065, supplied by HipSci). iPSCs were maintained at 37 °C and in 5% CO_2_ in mTeSR or mTeSR plus medium (Stem Cell Technologies, #85850 and #100-0276) on Geltrex-coated (Thermo Fisher Scientific, #A1413302) cell culture plates. Cells were split when confluent using ReLeSR (Stem Cell Technologies, #05873).

## METHODS DETAILS

### Plasmids

Sequences of β23 and Mito-GFP were previously described.^42,116^ Mutant ornithine transcarbamylase aa 30-114 deleted (OTCΔ) was a gift from Nicholas Hoogenraad (Addgene plasmid # 71878; http://n2t.net/addgene:71878; RRID: Addgene_71878).^30^ Mitochondrially targeted β23 (Mito-β) was created by modifying the β-sheet protein with a mitochondrial targeting sequence (MTS) derived from Neurospora crassa ATP synthase subunit 9 (Su9)^116^ at the N-terminus. For yeast experiments, the original MTS of OTCΔ was replaced with the Su9-derived MTS to enhance targeting efficiency. For human cell experiments, the original OTCΔ sequence was used. A myc-epitope tag was added to all constructs for detection. The Mito-β, OTCΔ, and GFP sequences were cloned into different plasmids for expression in yeast and human cells. The pRS416GAL1 (galactose-inducible) and pRS316GPD and pRS413GPD (constitutive) plasmids were used for yeast expression.^117,118^

For human cell expression the Mito-β and Mito-GFP sequences were cloned into pcDNA3.1(+) plasmid (Invitrogen). To generate a doxycycline (Dox)-inducible Mito-β construct, Mito-β was cloned into pCW57.1-Cas9 all-in-one Tet-on plasmid.^119^ Dox was used at 0.5 µg/mL. In addition, Mito-β and Mito-GFP were cloned into pLenti plasmid^120^ for expression in iNeurons. Lenti-EF1a-dCas9-VPR-Puro was a gift from Kristen Brennand (Addgene plasmid # 99373; http://n2t.net/addgene:99373; RRID: Addgene_99373). pCW-Cas9 was a gift from Eric Lander and David Sabatini (Addgene plasmid # 50661; http://n2t.net/addgene:50661; RRID: Addgene_50661).

For live cell imaging, we tagged Mito-β with the fluorescent protein mScarlet (Mito-mScarlet-β).^121^ pmScarlet-i_C1 was a gift from Dorus Gadella (Addgene plasmid # 85044; http://n2t.net/addgene:85044; RRID: Addgene_85044).

### Yeast transformation

To prepare competent yeast cells, 8 mL of exponentially growing cells were pelleted at 1,000 x g for 3 min, washed with water, pelleted, washed with SORB buffer (100 mM lithium acetate, 10 mM Tris-HCl pH 8.0, 1 mM EDTA pH 8.0, 1 M sorbitol) and pelleted again. The cells were resuspended in 225 µL SORB buffer. 25 µL herring sperm DNA (Invitrogen) was denatured at 95 °C for 5 min and added to the cells. Cells were aliquoted and stored at −80 °C.

For transformation of yeast cells, 200 ng DNA and 300 µL PEG buffer (100 mM lithium acetate, 10 mM Tris-HCl pH 8.0, 1 mM EDTA pH 8.0, 40% PEG 3350) were added to 50 µL competent cells. Cells were incubated at 30 °C for 30 min, 40 µL DMSO was added and incubated at 42 °C for 15 min. Cells were then pelleted, washed with water, pelleted again, resuspended in water and plated on selection plates.

### Generation of yeast deletion strains

Gene deletions in yeast were performed as previously described with modifications.^122^ A kanamycin resistance gene was amplified from the plasmid pFA6-kanMX4 with the addition of 50 base pairs of homologous regions corresponding to the genomic sequence immediately before and after the gene of interest. Yeast was transformed with the respective PCR product as described above with the difference that cells were grown in YPD medium overnight before plating on YPD plates with G418 (Thermo Fisher) selection. Positive clones were transferred to new selection plates. Successful integration of DNA by homologous recombination for gene deletions was verified by colony PCR. Primers were designed to bind within the deleted gene and therefore amplify only in control cells and not in cells with the desired gene deletion. 100 µL exponentially growing cells were pelleted and resuspended in 100 µL 20 mM NaOH, boiled at 99 °C for 10 min and frozen at - 20 °C. PCR was performed with1 µL of the lysate using MyFi polymerase after manufacturer’s protocol (biocat).

### Differentiation of iPSCs to neurons (iNeurons)

Neural progenitor cells (NPCs) were generated using the STEMdif SMADi Neural Induction Kit (Stem Cell Technologies, #08581) according to the monolayer protocol. NPCs were frozen in STEMdif Neural Induction Medium + SMADi with 10% DMSO.

Forebrain-type neurons were generated from the NPCs described above. NPCs were thawed in STEMdiff Neural Induction Medium + SMADi on 0.01% poly-L-ornithine/10-20 µg/ml laminin-coated (Merck #P4957 and #L2020) cell culture plates and differentiated using the STEMdiff Forebrain Neuron Differentiation Kit (Stem Cell Technologies, #08600) followed by the STEMdiff Forebrain Neuron Maturation Kit (Stem Cells Technologies, #08605). STEMdiff Forebrain Neuron Maturation medium was supplemented with 5 µM 5-F 5-fluorouracil/uridine (Merck #F6627, #U3750) for 6 days to stop the growth of undifferentiated cells. After 8 days in STEMdiff Forebrain Neuron Maturation Medium, iPSC-derived neurons were maintained in Neurobasal Plus Medium (Thermo Fisher Scientific, #A3582901) supplemented with B-27 Plus Supplement (Thermo Fisher Scientific, #A3582801), 0.5 mM GlutaMAX (Thermo Fisher Scientific, #35050061) and 100 U/ml penicillin, 100 µI/ml streptomycin sulfate (Thermo Fisher Scientific, #15140122).

### Lentiviral vector generation and transduction

Lentiviral vectors were generated to transduce HEK 293T cells with the goal of creating stable cell lines. Lenti-X HEK 293T cells (Takara) were grown in 10 cm dishes and transfected using Lipofectamine 3000 with the plasmids: pMD2.G (1.4 µg), psPAX2 (4.6 µg) and the plasmid of interest (6.5 µg). 2 days after transfection, the virus-containing medium was collected and passed through a 0.2 µm filter to remove cells. Lenti-X Concentrator (Takara) was added, the mixture was incubated for 1-2 h at 4 °C and centrifuged at 1500 x g for 45 min. The pellet was resuspended in PBS, and added to cells or was frozen at −80 °C.

HEK 293T cells were transduced with virus and 1 µg/mL polybrene and incubated for 3 days before selection with 2 µg/mL puromycin (Thermo Fisher). Upon completion of selection, assessed by the death of non-transduced cells in a separate well, transduced cells were sorted into single cells by FACS (BD FACS Aria III). The resulting monoclonal cell lines were verified by testing the expression of the insert or, in the case of shRNA constructs, the knockdown efficiency. “A detailed protocol is available here: dx.doi.org/10.17504/protocols.io.eq2lyxz3wgx9/v1”

### Knockdown of HSP60, LONP1 and HSP10

Stable monoclonal cell lines carrying IPTG-inducible HSP60 and scrambled shRNA (Vector Builder) were generated by lentiviral transduction. The shRNA was expressed by addition of 200 µM IPTG for 6 days.

LONP1 and HSP10 knockdown in HEK 293T cells was achieved by transfecting LONP1 or HSP10 siRNA, with non-targeted siRNA as control (Dharmacon, smartpool). 1.25 µL siRNA was used from a 20 µM stock, resulting in a final concentration of 41.7 nM after addition to cells. The siRNA was added to 50 µL Opti-MEM (Thermo Fisher) and 2 µL DharmaFECT 1 transfection reagent (Dharmacon) was added to another 50 µL Opti-MEM. Both were mixed after 5 min of separate incubation. The mixture was incubated for 15 min and added to 80% confluent cells in one well of a 24-well plate. Cells were incubated for 3 days before analysis.

Alternatively, siRNA KD was achieved by using Lipofectamine RNAiMAX (Thermo Fisher). 1.25 µL siRNA was used from a 20 µM stock and added to 25 µL Opti-MEM. 1 µL RNAiMAX was added to another 25 µL Opti-MEM. Both were mixed and incubated for 15 min. The mixture was added to 75,000 suspended cells in 450 µL DMEM in one well of a 48-well plate. Cells were incubated for 4 days before analysis.

Double knockdowns were achieved by combining HSP60 shRNA for 6 days and LONP1 siRNA for 3 days (DharmaFECT 1 transfection), with a longer time for HSP60 shRNA to allow for comparable knockdown efficiencies.

### Cell harvest and lysis for SDS-PAGE

Yeast cells: 1-2 OD600 cells grown in synthetic dropout medium were pelleted at 5,000g for 1 min, resuspended in 100 µL 0.1 M NaOH for 5 min, and pelleted at 10,000 x g for 1 min. The pellet was resuspended in lithium dodecyl sulfate (LDS) sample buffer (Thermo Fisher) and heated at 95 °C for 3 min. Cell debris was pelleted at 10,000 x g for 1 min prior to sodium dodecyl sulfate (SDS)-PAGE.

HEK 293T cells: Cells from one confluent well of a 12-well plate were trypsinized, pelleted at 500 x g for 1 min at 4 °C, washed with PBS, pelleted again, and lysed in 50 µL RIPA lysis and extraction buffer (Thermo) supplemented with protease inhibitor cocktail (Roche) and benzonase for at least 15 min on ice. Protein concentration was determined by Bradford assay (Bio-Rad).

### Solubility assay

Yeast cells: yeast was grown in galactose medium for 24 h to induce expression of Mito-GFP, OTCΔ or Mito-β. 10 OD of cells were pelleted at 3,000 x g for 3 min, washed with water, pelleted again at 3,000 x g for 3 min, and lysed with glass beads in lysis buffer (25 mM Tris-HCl, pH 7.4, 150 mM NaCl, 1 mM EDTA, 5% glycerol, 1% Triton X-100, protease inhibitor cocktail [Roche]). The lysate was cleared by centrifugation at 2,000g for 5 min at 4 °C.

HEK 293T cells: Cells were grown in 12-well plates and transfected with Mito-GFP, OTCΔ or Mito-β expression plasmids by Lipofectamine 3000. 24 h after transfection, cells were trypsinized, pelleted at 500 x g for 1 min, washed with PBS, pelleted again and lysed in RIPA buffer (Thermo) with protease inhibitor cocktail and benzonase for at least 15 min on ice. HEK 293T cell lysates were not cleared by centrifugation.

The supernatant was divided into two equal parts; one half was denatured and used as total and the other half was centrifuged at 16,000 x g for 10 min at 4 °C. The supernatant was collected and denatured, the pellet was washed with lysis buffer and centrifuged at 16,000 x g for 10 min at 4 °C. The pellet was solubilized and denatured in LDS sample buffer (Thermo Fisher). The total, soluble and pellet fractions were analyzed by SDS-PAGE.

### SDS-PAGE and immunoblotting

Proteins were separated on NuPAGE 4-12% Bis-Tris protein gels (Invitrogen) using NuPAGE MES running buffer (Invitrogen) at 200 V. Proteins were transferred to PVDF membranes by semi-dry transfer (48 mM Tris, 39 mM glycine, 1.3 mM SDS, 20% methanol) at 100 mA per membrane for 75 min. Membranes were blocked in 5% skim milk in TBST buffer (20 mM Tris-HCl pH 7.6, 150 mM NaCl, 0.1% Tween-20), incubated in primary antibody overnight at 4 °C in 5% skim milk in TBST buffer, washed 3-times with TBST, incubated in secondary HRP-antibody for at least 1 h at room temperature (RT), and washed 3-times for 5 min with TBST. Membranes were developed with Luminata/Immobilon HRP substrate (Merck) and imaged with ImageQuant800 (Amersham). Protein bands were quantified with AIDA Image Analyzer v.4.27 (Elysia Raytest). Images were processed with Fiji 1.53q.

### Aggregate interactome analysis

Yeast cells containing galactose-inducible Mito-β, Mito-GFP and EV were grown overnight in raffinose medium with heavy (Lys_8_), medium (Lys_4_) and light (Lys_0_) lysine for SILAC labelling, respectively. Cells were then grown in SILAC-galactose medium for 24 h to induce expression. 50 OD of exponentially growing cells were harvested and resuspended in lysis buffer (25 mM Tris-HCl, pH 7.4, 150 mM NaCl, 1 mM EDTA, 5% glycerol, 1% Triton X-100, protease inhibitor cocktail (Roche)). Cells were lysed with glass beads using a FastPrep-24 homogenizer with a CoolPrep adapter (MP Biomedicals). Lysates were cleared by centrifugation at 2,000 x g for 5 min at 4 °C. 50 µL anti-myc µBeads suspension (Miltenyi Biotech) was added to 1.5 mg protein in 1 mL lysis buffer and slowly rotated at 4 °C for 1 h. Lysates were applied to µ-columns (Miltenyi Biotech) and washed 4-times with 250 µL lysis buffer and once with PBS before eluting with 70 µL hot LDS sample buffer. Eluates were combined and processed using the Filter Aided Sample Prep (FASP) method (see below).

### Insoluble proteome of yeast cells

Yeast cells containing galactose-inducible Mito-β and EV control cells were grown overnight in raffinose medium in heavy (Lys_8_) and light (Lys_0_) lysine for SILAC-labelling, respectively. Afterwards, cells were grown for 24 h in SILAC galactose medium to induce expression. 50 OD exponentially growing cells were pelleted at 3,000g for 3 min, washed with 1 mL water, pelleted again at 3,000 x g for 1 min and lysed with glass beads in lysis buffer (25 mM Tris-HCl, pH 7.4, 150 mM NaCl, 1 mM EDTA, 5% glycerol, 0.5% Triton X-100, protease inhibitor cocktail [Roche]). Lysate was cleared by centrifugation at 800 x g for 5 min at 4 °C. For pellet fractions, 1 mg protein lysate in 500 µL was centrifuged at 20,000 x g for 20 min at 4 °C, washed with 0.5% Triton X-100 lysis buffer and centrifuged again at 20,000 x g for 20 min at 4 °C. LDS sample buffer (Thermo Fisher) was added to the pellet, followed by sonication in a Bioruptor (Diagenode) on high until dissolved. Pellet fractions were combined with 200 µg of corresponding total fraction (5x more pellet than total to increase detection of low abundant pelleted proteins). Peptides were generated with FASP and analyzed by mass spectrometry. The insoluble fraction of proteins, defined as amount in pellet divided by amount in total, were corrected for higher pellet concentration.

### Insoluble proteome of HEK 293T cells

HEK 293T cells with Dox-inducible Mito-β and wildtype (WT) cells were grown with Dox for 3 days. Cells from a confluent 6 cm dish were harvested by trypsinization, pelleted at 500 x g for 1 min at 4 °C, washed with PBS, pelleted again at 500 x g for 1 min at 4 °C, and lysed in 100 µL 1% Triton X-100 in PBS with protease inhibitor cocktail and benzonase for at least 15 min on ice. Protein concentration was determined by Bradford assay. For the pellet fraction, 500 µg protein was diluted in 250 µL 1% Triton X-100 lysis buffer, centrifuged at 20,000 x g for 20 min at 4 °C, washed with 1% Triton X-100 lysis buffer and centrifuged again at 20,000 x g for 20 min at 4 °C. The pellet was solubilized in 50 µL lysis buffer (iST, Preomics) and sonicated 5-times (30 s on, 15 s off) in a Bioruptor (Diagenode) on high setting. For total fractions, 50 µg protein was denatured in 50 µL lysis buffer. Total and pellet fractions were incubated at 95 °C for 10 min, peptides were generated with the iST-Kit (Preomics) and analyzed by label-free mass spectrometry. Insoluble fractions of proteins, defined as amount in pellet divided by amount in total, were corrected for higher pellet concentration.

### Pulse-labeling of newly-synthesized protein in HEK 293T cells

HEK 293T cells with Dox-inducible Mito-β or EV were grown with 0.5 µg/mL Dox in standard light medium (L) for 16 h, followed by labeling with heavy lysine (Lys_8_) and arginine (Arg_10_) (H) for 4 h. Cells were separated into pellet and total fractions as before. Lysate from cells labeled with medium lysine (Lys_4_) and arginine (Arg_6_) (M) for at least 5 days was added to the pellet and total fractions as spike-in prior to peptide generation and SILAC MS analysis. To determine the insoluble fraction of H-labeled proteins, the H/M ratio of the pellet fraction was divided by 10 and by the H/M ratio of the total fraction. Similarly, for unlabeled (L) proteins, the L/M ratio of the pellet fraction was divided by 10 and by the L/M ratio of the total fraction.

### Isolation of mitochondria and mitoribosome separation by sucrose gradient

Isolation of mitochondria from HEK 293T cells was performed as described previously with modifications.^123^ Cells from a 10 cm dish were harvested by trypsinization and frozen at −20 °C as a dry pellet. To swell the cells, the pellet was resuspended in 1 mL hypotonic T-K-Mg buffer (10 mM Tris-HCl, pH 7.4; 10 mM KCl; 0.5 mM MgCl_2_) per 0.15 g of pellet weight. Cells were incubated on ice for 5 min before lysis with a Dounce homogenizer with a tight-fitting Teflon pestle (2 ml capacity) with 30 strokes. Sucrose buffer (10 mM Tris-HCL, pH 7.4; 10 mM KCl; 0.5 mM MgCl_2_; 1 M sucrose) equal to 1/3 of the volume of the lysate was added immediately after lysis to make the lysate isotonic. Cell debris was removed from the lysate by centrifugation twice at 1200 x g for 3 min at 4 °C. The supernatant was centrifuged at 15,000 x g for 5 min at 4 °C to pellet the crude mitochondrial fraction. The pellet was resuspended in sucrose buffer (10 mM Tris-HCL, pH 7.4; 10 mM KCl; 0.5 mM MgCl_2_; 0.25 M sucrose) and centrifuged again at 15,000 x g for 5 min at 4 °C. The pellet was resuspended in 100 µL mitochondrial lysis buffer (10 mM Tris-HCl pH 7.5, 100 mM KCl, 20 mM MgCl_2_, 260 mM sucrose, 1% Triton X-100 (w/v), 1x protease inhibitor, 1x SUPERaseIn RNA inhibitor 0.1 U/µL) and incubated on ice for 30 min. During incubation, the protein concentration of the lysate was measured by Bradford assay. 50 µg was saved as total and 500 µg protein in 100 µL lysis buffer was centrifuged at 20,000 x g for 20 min at 4 °C to remove insoluble material. The supernatant was stored on ice.

The separation of mitoribosomes of HEK 293T cells by sucrose gradient was performed as described previously with modifications.^87^ 6.25 mL of ice-cold 10% sucrose buffer (20 mM Tris-HCl, pH 7.5; 100 mM KCl, 20 mM MgCl_2_, 10% or 30% sucrose (w/v)) was added to an ultracentrifugation tube (open-top poly clear (14×89 mm), Seton) and underlaid with 6.25 mL of 30% sucrose gradient buffer. A linear gradient was formed using a gradient maker (Biocomp). The weight of the sucrose gradients was balanced and the prepared mitochondrial lysates were loaded. Tubes were centrifuged in a SW41Ti rotor at 79,000 x g for 15 h at 4 °C. The gradient was fractionated into 12 fractions using a piston fractionator (Biocomp), with the last fraction containing the remaining buffer with the pellet.

Proteins in gradient fractions were precipitated by adding 250 µL 50% trichloroacetic acid to 1 mL of sample. Samples were vortexed and incubated for 30 min on ice. Samples were centrifuged at 10,000 x g for 10 min at 4 °C and the supernatant was discarded. Pellets were washed twice by adding 500 µL ice-cold acetone, vortexing and centrifuging at 10,000 x g for 10 min at 4 °C. The supernatant was removed and the pellets were allowed to dry before the addition of lysis buffer (Preomics). Samples were sonicated in a Bioruptor (Diagenode) until the pellets dissolved and heated at 95 °C for 10 min. Peptides for LC-MS/MS were generated using iST kits (Preomics) according to the manufacturer’s protocol.

### Sample preparation for mass spectrometry

Peptides for LC-MS/MS were generated using the Filter Aided Sample Prep (FASP) method.^124,125^ All centrifugation steps until peptide solubilization were performed at 14,000 x g for 15 min at RT. Samples were diluted with urea buffer 1:4 (UA) (8 M urea, 0.1 M TRIS pH 8.5), added to an Amicon Ultra 0.5 mL 10 kDa filter (Millipore) and centrifuged. Then, 200 µL UA was added to the sample and centrifuged. This process was repeated 2-times more. 200 µL of 10 mM DTT in UA was added, the sample incubated for 45 min at RT and centrifuged. 100 µL of 55 mM iodoacetamide in UA was added, followed by incubation for 30 min at RT in the dark and centrifugation. 200 µL UA was added to the sample and centrifuged. This process was repeated 2-times more. The samples were then washed twice with 200 µL of 40 mM ammonium bicarbonate (NH_4_HCO_3_). 40 µL protease solution (30 ng/µL) was added. Yeast proteins were digested with lysyl endopeptidase (Wako) and human proteins were digested with trypsin (Roche) by incubation at 37 °C overnight. Peptides were eluted by centrifugation, followed by the addition of 50 µL 40 mM NH_4_HCO_3_ and centrifugation. Another 50 µL of 40 mM NH_4_HCO_3_ was added and samples centrifuged. 5 µL of 25% trifluoracetic acid (TFA) was added and the peptides were completely dried in a vacuum centrifuge at 45 °C. Pellets were solubilized in 30 µL 0.1% TFA.

To desalt the peptides, home-made microcolumns containing C18 Empore disks were washed by adding 50 µL acetonitrile (ACN) and centrifugation at 1,000 x g for 3 min. 50 µL 0.1% TFA was added to the column, followed by centrifugation as before. Peptides were added and columns centrifuged at 240 x g for 5 min or until liquid had passed the column. The flowthrough was reapplied and columns centrifuged as before. 100 µL 0.1% TFA was added to the columns, followed by centrifugation at,1,000 x g for 5 min. Peptides were eluted with 30 µL 70% ACN 1% formic acid (FA) at 240 x g for 5 min into a new tube. Eluates were dried in a vacuum centrifuge at 45 °C, resuspended in 10 µL 5% FA and sonicated in an ultrasonic water bath for 5 min. Alternatively, peptides for LC-MS/MS were generated using iST kits (Preomics) according to the manufacturer’s protocol.

### Mass spectrometry

Mass spectrometry was performed on a Q-Exactive HF mass spectrometer with a nanospray ion source and a nanoHPLC Proxeon EASY-nLC 1200 system (Thermo Fisher). Reversed-phase separation was accomplished using a home-made C18 AQ phase (1.9 μm, Dr. Maisch) packed silica capillary (75 μm inner diameter × 30 cm length). 0.1% FA in H_2_O was used as mobile phase A, and 0.1% FA in 80% ACN was used as mobile phase B. Samples were separated with a linear gradient from 5 to 35% buffer B in 120 min at a flow rate of 250 nL/min. The Q-Exactive was operated in data-dependent mode with survey scans acquired at a mass range from 300 to 1650 m/z and a resolution setting of 60,000 (FWHM). The MS1 AGC target was set to 3e6 with a maximum injection time of 50 ms. Up to ten of the most abundant ions from the survey scan were selected and fragmented by higher energy collisional dissociation with an energy setting of 29. MS/MS spectra were acquired with a resolution of 15,000 (FWHM), a maximum injection time of 50 ms, a precursor isolation width of 1.2 Da, and a target value of 1e2.

### Mass spectrometry database search

The acquired mass spectrometric raw data were processed using MaxQuant version 1.6.3.4 (https://www.maxquant.org/). MS/MS spectra were searched against human proteins (Uniprot version 20201109 (insoluble proteome Mito-β), 20211108 (insoluble proteome KDs)) or the yeast proteome (Uniprot version 20181109 (interactome Mito-β and OTCΔ), 20201104 (insoluble proteome of Mito-β and LONP1 deletion/KD), 20221026 (Total proteome)). As search parameters, trypsin (LysC for yeast) allowing for cleavage N-terminal to proline was chosen as enzyme specificity; for SILAC-labelled samples, Arg_6_, Lys_4_ were selected for medium-labels and Arg_10_, Lys_8_ for heavy labels (only Lys for yeast experiments); cysteine carbamidomethyl (C) was set as fixed modification, while protein N-terminal acetylation and methionine oxidation were selected as variable modifications. A maximum of two missed cleavages and three labels were allowed. Proteins and peptides (minimal seven amino acids) were identified at a false discovery rate of 1% using a target-decoy approach with a reversed database. The “re-quantify” option was enabled for protein quantification. Label-free quantitation was performed with default settings and "match between runs" functionality enabled. Proteins were included in the analysis if they were identified in at least 2 repeats per condition.

### Tetrazolium overlay assay

The frequency of respiratory-deficient yeast colonies was quantified by the tetrazolium assay, which colors normal colonies deep red due to the reaction of tetrazolium to formazan by mitochondrial dehydrogenases, while petite colonies remain white, as previously described with modifications.^72^ Yeast cells were grown as single colonies on glucose plates (galactose for inducible strains). One colony was picked, dissolved in water and 1/1,000 of it was plated on a second plate. After colonies formed, 1.5% low melting agar was dissolved in medium and cooled to 40-50 °C before 0.2% 2,3,5-triphenyltetrazolium chloride (Sigma-Aldrich) was added and poured over the plates. Plates were imaged after 3 h at 30 °C or until control colonies turned red.

### Staining of mtDNA in yeast

Staining of mtDNA was performed in live yeast cells by incubating cells in medium containing 1 µg/µL DAPI for 15 min at RT. µ-Slides (Ibidi) were coated with Concanavalin A for 5 min and washed with PBS. After incubation, cells were pelleted by centrifugation at 3,000 x g for 1 min, washed with PBS, pelleted again, resuspended in medium and applied to µ-slides. Slides were incubated at RT for 10 min to allow cells to attach and washed with medium. Cells were imaged under a fluorescence microscope at RT.

### Staining of mtDNA in HEK 293T cells

Staining of mtDNA was performed as previously described with modifications.^81^ HEK 293T cells were grown in 12-well plates with 0.5 µg/mL Dox to induce Mito-β expression, and transferred to poly-L-lysine-coated 8-well microslides (Ibidi) one day before imaging. mtDNA was stained by incubation in SybrGold (Invitrogen) diluted 1:10,000 in DMEM for 5 min at 37 °C. Cells were washed with PBS and imaged under a fluorescence microscope at RT in DMEM.

### mtDNA quantification by qPCR

Isolation of gDNA and mtDNA was performed using a NucleoSpin Tissue Mini kit for DNA from cells and tissue (Macherey-Nagel). 10 ng total DNA was used for the PCR reaction with PowerUp SYBR Green Master Mix (Applied Biosystems). 0.5 µM primers were used against the mitochondrial 16S rRNA gene (16S F: cgaaaggacaagagaaataagg, 16S R: ctgtaaagttttaagttttatgcg) and the β-globin gene as a reference (βG F: caacttcatccacgttcacc, βG R: gaagagccaaggacaggtac).^126^ Relative mitochondrial DNA copy number was determined by the ΔΔct method. ct(16S) and ct(βG) are the cycle threshold values of the qPCR reaction of 16S rRNA and the β-globin mRNA, respectively.

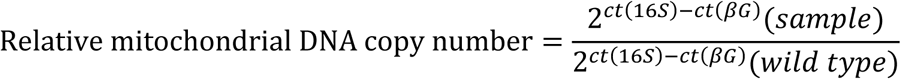

### RNA quantification by qPCR

Total RNA was isolated from cells using the RNeasy Mini Plus Kit (Qiagen, #74134). For cDNA synthesis, 1 µg RNA in 10 µL was combined with 1 µL random hexamer primer (0.2 µg/µL) (Thermo Fisher), incubated at 70 °C for 3 min and cooled on ice. 1 µL dNTPs (10 mM), 2 µL RNAse-free water, 1 µL RNasin ribonuclease inhibitors (Promega), 4 µL 5x reverse transcriptase buffer (Thermo Fisher) and 1 µL Maxima reverse transcriptase (Thermo Fisher) were added. The mixture was incubated in a thermocycler with the following settings: 10 min 25 °C, 30 min 50 °C, 10 min 55 °C, 10 min 60 °C, 10 min 65 °C, 5 min 85 °C and 1 min 25 °C. 10 ng of cDNA was used for quantitative real-time PCR using PowerUp™ SYBR™ Green Master Mix (Applied Biosystems #A25776) on a QuantStudio 1 Real-Time PCR System (Applied Biosystems). Ct values were measured and fold changes were calculated using the ΔΔct method with the RPL27 gene as reference. The following primers were used:

RPL27 forward: 5’-GCTGGAATTGACCGCTACC-3’ and reverse: 5’-TCTCTGAAGACATCCTTATTGACG-3’; 12S forward: 5’-CCTCACCACCTCTTGCTCAG-3’ and reverse: 5’-TTACGTGGGTACTTGCGCTT-3’;

16S forward: 5’-ACCCTCACTGTCAACCCAAC-3’ and reverse: 5’-TGGTAAACAGGCGGGGTAAG-3’; COX2 forward: 5’-ACGCATCCTTTACATAACAGAC-3’ and reverse: 5’-GCCAATTGATTTGATGGTAAGG-3’.

### Transcriptomics

RNA isolation was performed using TRI reagent (Sigma-Aldrich, #T9424) according to the manufacturer’s instructions. Yeast cells were grown in glucose medium overnight and pelleted by centrifugation or transferred to glycerol medium for 5 h and then pelleted. 25 OD of cells were resuspended in 1 mL of TRI reagent and lysed with glass beads using a FastPrep-24 homogenizer with a CoolPrep adapter (MP Biomedicals). Samples were incubated at RT for 5 min to dissociate nucleo-protein complexes. 0.2 mL of chloroform was added per 1 mL of TRI reagent used for cell lysis. Samples were shaken vigorously for 15 s, incubated at RT for 10 min, centrifuged at 12,000 x g for 10 min at 4 °C. The upper aqueous phase was transferred to a fresh tube and supplemented with isopropanol (0.5 mL per 1 mL of TRI reagent used). The sample was mixed by inversion and incubated for 5-10 min at RT. The RNA pellet was washed with 1 mL of 75% ethanol by centrifugation at 10,000 x g for 10 min at 4 °C. The pellet was air dried and dissolved in water. RNA concentration was measured using Nanodrop 1000. Library preparation was performed using TruSeq Stranded mRNA according to the manufacturer’s protocol. Samples were sequenced on an Illumina NextSeq 500.

RNA-seq reads were trimmed with TrimGalore (http://www.bioinformatics.babraham.ac.uk/projects/trim_galore/, v0.4.0) and mapped to the yeast annotation using TopHat2^127^ (v2.0.13) with the options mate-inner-dist, mate-std-dev and library-type (fr-firststrand). The distance between read mates (mate-inner-dist and mate-std-dev) were assessed individually using Picard tools [http://broadinstitute.github.io/picard.] (v1.1.21). After mapping of the RNA-seq reads from all samples, the reads that mapped uniquely to the genome were counted using featureCounts^128^ with the following options: -Q 10 -p -B -C -s 2. DESeq2^129^ was used for differential expression analysis of all replicates for each condition.

### Yeast spot assay

Yeast cells containing plasmids for expression of Mito-GFP, OTCΔ or Mito-β were grown overnight to stationary phase in glucose medium for constitutive expression and in raffinose medium for galactose-induction. 1 OD of cells was pelleted at 3,000 x g for 3 min and resuspended in 100 µL water. 25 µL yeast suspension was added to 225 µL water in the first wells of a 96-well plate. 50 µL of the first well was transferred to the second well containing 200 µL. Such 1:5 dilution series steps were continued 4 more times. Cells were transferred to glucose, galactose, and glycerol plates with a stamp and incubated until colonies appeared in the most diluted spot.

### Human cell viability assay

HEK 293T WT cells and cells with Dox-inducible Mito-β expression were grown in 12 well plates with 0.5 µg/mL Dox in DMEM with glucose or galactose for up to 7 days without letting them reach full confluency. The reaction of 3-(4,5-dimethylthiazol-2-yl)-2,5-diphenyltetrazolium bromide (MTT) to formazan by respiratory active cells was used as a readout for cell viability. Cells were incubated in 0.5 mg/mL MTT in DMEM for 1 h at 37 °C. The reaction was stopped by adding 0.5 volume of stop solution (40% N,N-Dimethylformamide 2% glacial acetic acid, 16% SDS) and the plates were shaken until formazan crystals dissolved. Formazan was quantified by measuring absorbance at 570 nm in a plate reader spectrometer (CLARIOstar BMG Labtech).

### Quantification of mitochondrial translation

#### Yeast cells

A yeast strain carrying mt-encoded superfolder GFP^61^ (GFP^mtDNA^) was transformed with galactose-inducible Mito-β, OTCΔ, and EV control constructs. Cultures were grown in galactose medium for 24 h and GFP quantified by immunoblotting with anti-GFP antibody.

#### HEK 293T cells

Newly-synthesized mitochondrial proteins were quantified by L-homopropargylglycine (HPG) incorporation (Invitrogen, Click-iT™ HPG Alexa Fluor™ 488 Protein Synthesis Assay Kit) and fluorescent labeling by Click chemistry, as previously described with modifications.^64^ Cells were grown on 8-well slides (Ibidi) and incubated with HPG and the cytosolic translation inhibitor emetine (50 µg/mL) in methionine-free medium for 1 h. After labeling, the slides were placed on ice and incubated in ice-cold buffer A (10 mM HEPES/KOH pH 7.5; 10 mM NaCl; 5 mM MgCl_2_; 300 mM sucrose) supplemented with 0.015% digitonin for 2 min. Cells were fixed after permeabilization in buffer A containing 4% formaldehyde for 10 min. Cells were washed with 3% BSA in PBS, incubated with 0.5% Triton X-100 in PBS for 10 min, and washed again with 3% BSA in PBS. Click reaction was performed as described in the manual of the Click-iT™ HPG Alexa Fluor™ 488 Protein Synthesis Assay. To visualize Mito-β and OTCΔ, cells were incubated in anti-myc or anti-OTC antibody overnight, washed with 3% BSA, incubated in secondary antibody for at least 1 h and washed twice with PBS. Cells were imaged under a fluorescence microscope (Zeiss LSM980 with Airyscan 2 module), and the mean fluorescence signal intensity in single cells was quantified using ImageJ software.

### Quantification of reactive oxygen species

Yeast cells were incubated with 5 µM CellROX for 1 h at 30 °C, pelleted at 3,000 x g for 1 min, washed with water, pelleted again and resuspended in water. The mean fluorescence intensity was quantified by flow cytometry (Attune NxT, Thermo Fisher).

### Iron quantification

100 OD of exponentially growing yeast was pelleted at 3,000 x g for 1 min and washed twice with water. Cells were incubated in 1 mL 0.1 M NaOH for 5 min, centrifuged at 5,000 x g for 1 min, lysed in lysis buffer (5% SDS, 200 mM Tris pH 6.8, 2% β-mercaptoethanol) and incubated at 98 °C for 5 min. Iron was quantified using a colorimetric bathophenanthrolinedisulfonic acid (BPS) assay.^130^ The reaction was performed in 100 µL consisting of 90 µL cell lysate in 33 mM ascorbic acid and 1 mM BPS. The absorbance was measured at 535 nm - 680 nm in a plate reader (CLARIOstar BMG Labtech).

### Immunofluorescence

#### Yeast

Yeast cells were cultured in glucose medium for 24 h and fixed in exponential phase with 4% formaldehyde solution for at least 1 h. Spheroplasts were prepared by incubation in 100 µg/mL Zymolase 100T (Roth) in 0.1 M potassium phosphate buffer pH 7.5 with 2 µL/mL β-mercaptoethanol for 1 h at 30 °C or until spheroplasting was complete, as judged by less reflective appearance under a phase-contrast light microscope. Spheroplasts were pelleted at 500 x g for 3 min, resuspended in PBS, applied to poly-L-lysine-coated coverslips and allowed to settle for 10 min.

#### iNeurons

Cells were grown on poly-L-lysine-coated coverslips, fixed for 10 min in 4% formaldehyde in PBS and washed with PBS.

#### Yeast and iNeurons

Cells were permeabilized with 0.1% Triton X-100 in PBS for 5 min followed by blocking with 3% BSA in PBS for at least 10 min. Primary antibody was applied overnight in 3% BSA in PBS and cells were washed 3-times with PBS. Secondary antibody was applied in 3% BSA in PBS for at least 1 h, followed by three PBS washes. Coverslips were mounted with mounting medium (DAKO) and imaged under a fluorescence microscope after drying.

### Visualization of protein aggregates

Protein aggregates were detected using the Proteostat protein aggregation kit (Enzo Life Sciences) as previously described.^84^ HEK 293T cells with scrambled shRNA and non-targeted siRNA control, LONP1 KD, HSP60 KD, combined LONP1 and HSP60 KD, or Dox-induced Mito-β expression were grown on poly-L-lysine-coated coverslips. Cells were fixed for 10 min in 4% formaldehyde in PBS. Cells were washed with PBS, permeabilized with 0.1% Triton X-100 in PBS for 5 min and washed with PBS. Aggregates were stained with Proteostat detection dye diluted 1/1000 in reaction buffer (Enzo Life Sciences) for 30 min in the dark. Cells were washed with PBS and incubated in 3% BSA in PBS for at least 10 min. To visualize mitochondria, cells were incubated with anti-TOM20 antibody overnight, washed with 3% BSA, incubated with secondary antibody for at least 1 h and washed twice with PBS. Coverslips were mounted with mounting medium (DAKO) and imaged under a fluorescence microscope after drying.

### Visualization of mitochondrial cristae

Mitochondrial cristae were visualized by exploiting the property of the mitochondrial dye MitoBright LT Green (Dojindo) to accumulate in the mitochondrial intermembrane space and cristae lumen. HeLa cells were transfected with Mito-mScarlet-β and transferred to 8-well µ-slides (Ibidi) the next day. After 2 days, the cells were treated with 0.1 µM MitoBright LT Green for 30 min and washed with DMEM. The cells were imaged live at RT by structured illumination microscopy (SIM) on a Zeiss Elyra PS.1 microscope with SIM module.

### DNA-PAINT super-resolution microscopy

DNA-points accumulation for imaging in nanoscale topography (DNA-PAINT)^131^ is a microscopy method in which fluorescence emitters are switched on and off stochastically, so that only a subset of fluorophores is on at the same time. This temporal separation of fluorescence emission allows a much higher precision of imaging than standard widefield microscopy, where all fluorophores are on at the same time. In DNA-PAINT, single-molecule on/off-switching is generated by transient binding of fluorophore-labeled DNA-imager strands to complementary DNA-docking strands, immobilized to the structure of interest in the sample. Imagers can bind repeatedly to their complementary docking strands and are therefore imaged in thousands of consecutive frames. A final DNA-PAINT image is reconstructed from all imaged frames by first fitting the most likely position of each single-molecule blinking event in each frame, and then overlaying all individual frames to obtain the final image. To apply DNA-PAINT in cells, labelling with primary antibodies as well as their cognate secondary binders, such as nanobodies or Fab fragments, is performed after fixation. For DNA-PAINT imaging, these secondary binders are covalently modified with single-stranded DNA docking strands, allowing them to be addressed by their complementary imagers.

#### Protein-DNA conjugation

Nanobodies were conjugated as described previously.^132^ Unconjugated nanobodies were thawed on ice, then a 20-fold molar excess of bifunctional DBCO-PEG4-maleimide (Jena Bioscience, CLK-A108P) was added and reacted for 2 h on ice. Unreacted linker was removed by buffer exchange to PBS using Amicon centrifugal filters (10,000 MWCO). The DBCO modified nanobodies were reacted with 5x molar excess of azide-functionalized docking strand DNA (5xR1, 5xR2, 7xR3, 7xR4 or 5xR5) overnight at 4 °C. Unconjugated protein and free DNA were removed by anion exchange chromatography using an ÄKTA Pure system equipped with a Resource Q 1-ml column. The AffiniPure Fab Fragment Bovine Anti-Goat IgG, Fc Fragment Specific (JAC805007008, Jackson ImmunoResearch) was reacted with 50x excess of TCEP and incubated for 30 min at RT. Then, the buffer was exchanged to PBS with 5 mM EDTA pH 7, using a 40k Zeba column (Thermo Fisher Scientific, 87766). The Fab fragment was reacted with 10x excess of DBCO-PEG4-maleimide and incubated for 1.5 h at 23 °C and 300 rpm. Unreacted linker was removed with a 40k Zeba column and the buffer was exchanged to PBS. DNA conjugation was performed by incubating the functionalized Fab with 10x excess of azide-DNA overnight at 25 °C.

#### Sample preparation for DNA-PAINT imaging

Hela or HEK 293T cells were seeded on µ-Slide 8 Well high slides (ibidi, 80806) and transfected with Mito-β the next day. After 1 or 2 days of expression, cells were fixed in a solution of 3% formaldehyde (Electron Microscopy Sciences, 15710) and 0.1% glutaraldehyde (Electron Microscopy Sciences, 16220) in PBS for 15 min. Cells were washed with PBS and quenched for 5 min with 0.2 M NH_4_Cl in PBS. Blocking and permeabilization was performed with a solution of 0.25% Triton X-100 (Carl Roth, 6683.1) and 3% BSA (Sigma-Aldrich, A4503) in PBS for 1.5 h. For all primary antibodies, cells were incubated with primary antibody in 0.1% Triton X-100/3% BSA overnight at 4 °C and washes 5 times with PBS. For all secondary binders, cells were incubated with secondary binder in nanobody incubation buffer at RT for 1.5 h. After 5 PBS washes, the samples were post-fixed with 4% PFA for 10 min and quenched again. After post-fixation of the anti-DNA/mkLCNb, the sample was permeabilized and blocked again. This procedure was repeated for every additional primary antibody and corresponding secondary binder. After staining and post-fixation, the sample was incubated with a 1:3 solution of gold nanoparticles (Cytodiagnostics, G-90-100) in PBS for 10 min and washed twice with PBS before proceeding to imaging.

#### DNA-PAINT imaging

Images were acquired using Exchange-PAINT, visualizing one target at a time.^133^ Prior to imaging, a fresh solution of 1 mM (±)-6-hydroxy-2,5,7,8-tetramethylchroman-2-carboxylic acid (Trolox) (Sigma-Aldrich, 238813)/ 10 µM 3,4-dihydroxybenzoic acid (Sigma-Aldrich, 37580)/ 2.5mM Pseudomonas protocatechuate 3,4-dioxygenase (Sigma-Aldrich, P8279) was prepared in buffer C (500 mM NaCl, 0.05% Tween 20 and 1 mM EDTA in PBS). The imager strand for the first target was added to this solution at a concentration of 50-500 pM. Before imaging the first target, the imager solution was incubated for 5 min and then replaced with fresh imager, after that the first acquisition round was started. Before introducing the imager specific for the next target of interest, the sample was washed with at least 2 ml of PBS until no residual signal from the previous imager solution was detected. Then, the next imager solution was added. All imager concentrations as well as acquisition times were adjusted according to the target density.

#### Microscope setup

Fluorescence imaging was carried out on an inverted microscope (Nikon Instruments, Eclipse Ti2) with the Perfect Focus System, applying an objective-type TIRF configuration equipped with an oil-immersion objective (Nikon Instruments, Apo SR TIRF ‘100, NA 1.49, Oil). A 560-nm laser (MPB Communications, 1 W) was used for excitation. The laser beam was passed through a cleanup filter (Chroma Technology, ZET561/10) and coupled into the microscope objective using a beam splitter (Chroma Technology, ZT561rdc). Fluorescence was spectrally filtered with an emission filter (Chroma Technology, ET600/50m and ET575lp) and imaged on an sCMOS camera (Andor, Zyla 4.2 Plus) without further magnification, resulting in an effective pixel size of 130 nm (after 2×2 binning). The transient binding of dye-labeled imager strands (R1: AGGAGGA-Cy3b, R2: GGTGGT-Cy3b, R3: GAGAGAG-Cy3b, R4: TGTGTGT-Cy3b, R5: GAAGAAG-Cy3b) to the docking strands (1xR1: TCCTCCT, 5xR1: TCCTCCTCCTCCTCCTCCT, 5xR2: ACCACCACCACCACCACCA, 7xR3: CTCTCTCTCTCTCTCTCTC, 7xR4: ACACACACACACACACACA, 5xR5: CTTCTTCTTCTTCTTCTTC) in the sample, was detected using a readout rate of 200 MHz. Images were acquired by choosing a region of interest with a size of 512 x 512 pixels. Raw microscopy data was acquired using μManager40 (Version 2.0.1). DNA-PAINT data was acquired using optical sectioning by highly inclined and laminated optical sheet (HILO) illumination.

#### DNA-PAINT analysis

Raw fluorescence data were subjected to super-resolution reconstruction using the Picasso software package (latest version available at https://github.com/jungmannlab/picasso).^131^ Drift correction was performed with a redundant cross-correlation and gold particles as fiducials.

#### Channel alignment

Alignment of subsequent imaging rounds was performed iteratively in Picasso, starting with a redundant cross-correlation and followed by gold fiducial alignment for cellular experiments.^131^

### Cryo-ET imaging

#### Sample preparation

Quantifoil EM grids (R1/2, Cu 200 mesh, square grids with holey carbon film for yeast cells and R1/2, Au 200 mesh grids with holey carbon film for HEK cells, QuantifoilMicroTools) were glow discharged using a plasma cleaner (PDC-3XG, Harrick) for 30 s. Yeast cells, grown in suspension culture until an OD_600_ of 0.6, were plunge frozen using a Vitrobot (Thermo Fisher) with the following settings: temperature, 30 °C, humidity, 80 %, blot time, 10 s, blot force, 10. 3.5 µl of yeast cell suspension were applied onto the carbon side of the glow discharged EM grids. Grids with yeast suspension were blotted from the back with Whatman paper and subsequently plunged into a liquid ethane/propane mixture.

HEK cells were trypsinized into single cells and afterwards sequentially plated onto gold EM-grids. The process was monitored in a light microscope to determine the confluency of HEK cells on the grid. HEK cells were added, until two to three cells occupied one grid square. HEK cells were cultivated for 24 h on the EM grids and subsequently plunge frozen into liquid ethane/propane using a Vitrobot with the following settings: temperature, 37 °C, 37 humidity, 80 %, blot time, 10 s, blot force, 10. The grids were blotted from the back and the front. The vitrified samples were transferred into cryo-EM boxes and stored in liquid nitrogen.

#### Cryo-FIB milling

EM grids were clipped under liquid nitrogen into autogrid sample carriers. For cryo-FIB milling, the samples were transferred into a Scios or a Quanta dual beam cryo-FIB/scanning electron microscopes (Thermo Fisher). To avoid damage of the sample by out of focus gallium ions, organometallic platinum was deposited onto the sample using a gas injection system (working distance: 10 mm, heating: 27 °C, time: 8 s). The grid was tilted to 18° with respect to the ion beam, and cellular material was removed from above and below the region of interest using gallium ions accelerated at 30 kV. For rough milling, currents of 0.3 nA and for fine milling currents of 50 pA were used. This resulted in thin cellular slices (lamellas), which had a thickness of 150 to 200 nm.

#### Data collection and reconstruction

The lamellas were transferred into a Titan Krios or a Tecnai Polara (Thermo Fisher) microscopes, both operated at 300 kV. Micrographs were collected using a 4 x 4 k K2 Summit (Gatan) direct electron detector operated in dose fractionation mode (0.2 s, 0.15 e^-^/Å^2^). Inelastically scattered electrons were removed by using a post column BioQuantum (Gatan) energy filter with a slit width of 20 eV. The tilt series were collected using SerialEM 3.7.0^134^ at pixel sizes of 0.352 and 0.439 nm. The collection was performed using a unidirectional acquisition scheme from −50° to + 60° with an angular increment of 2°. Individual frames were aligned using MotionCor2^135^ and tilt series was subsequently aligned based on fiducial-less patch tracking, and reconstructed using weighted back projection in IMOD 4.9.0.^136^

#### Tomogram segmentation

For segmenting cellular membranes, the tomograms were first filtered using the tomo_deconv filter (https://github.com/dtegunov/tom_deconv) to enhance the contrast. Then an automatic membrane tracing software package, TomoSegMemTV, ^137^ was used. The results were manually refined in Amira 6.2 (Thermo Fisher). Aggregates within mitochondria were segmented using the magic wand tool in Amira 6.2 by selecting areas of higher contrast.

### Quantification and statistical analysis

Statistical analysis of bar graph and box plot data was performed with Graphpad Prism 9 and analysis of proteomics data with Perseus. All statistical tests utilized are indicated within the respective figure legends. For multiple comparisons, One-way ANOVA with Dunnett’s or Tukey’s multiple comparison test was used. For multiple comparisons with normalized controls, Repeated-measures One-way ANOVA with Bonferroni′s or Dunnett′s multiple comparisons test was used. For multiple comparisons with normalized controls and unequal n, Mixed-effects analysis with Bonferroni′s or Dunnett′s multiple comparisons test was used. For comparison of one group to a reference value, One-sample Student’s t-tests was used.

## KEY RESOURCES TABLE

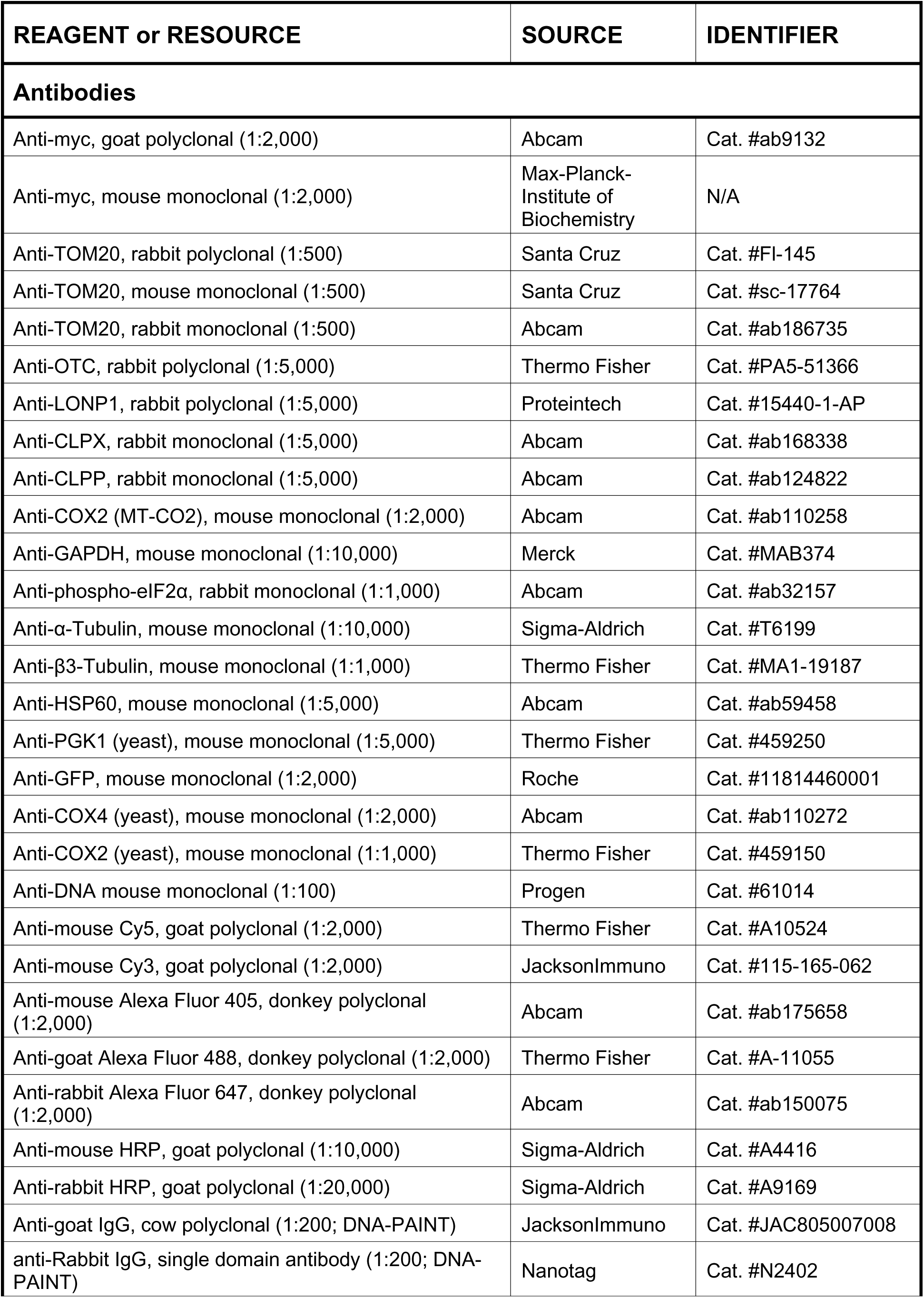
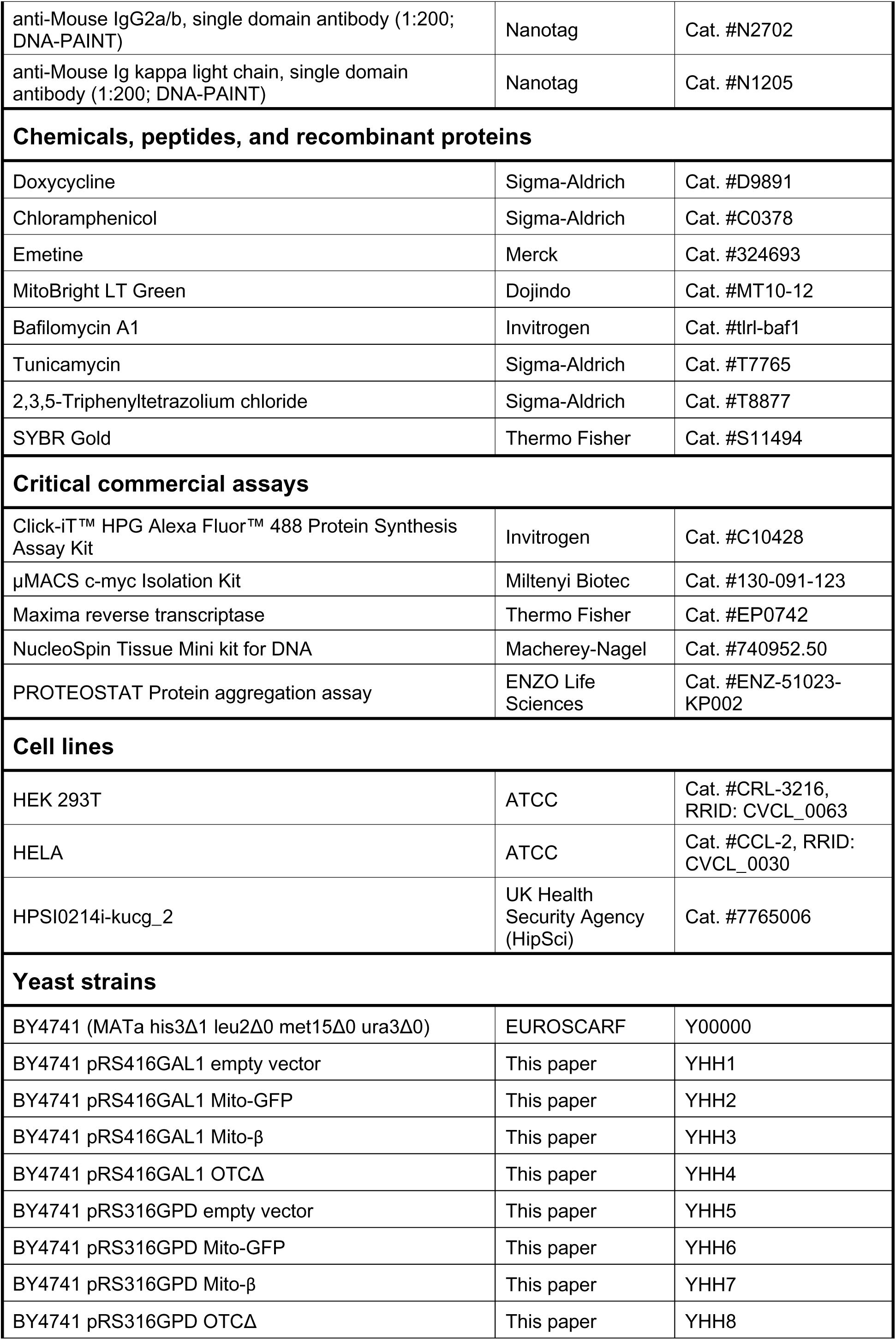
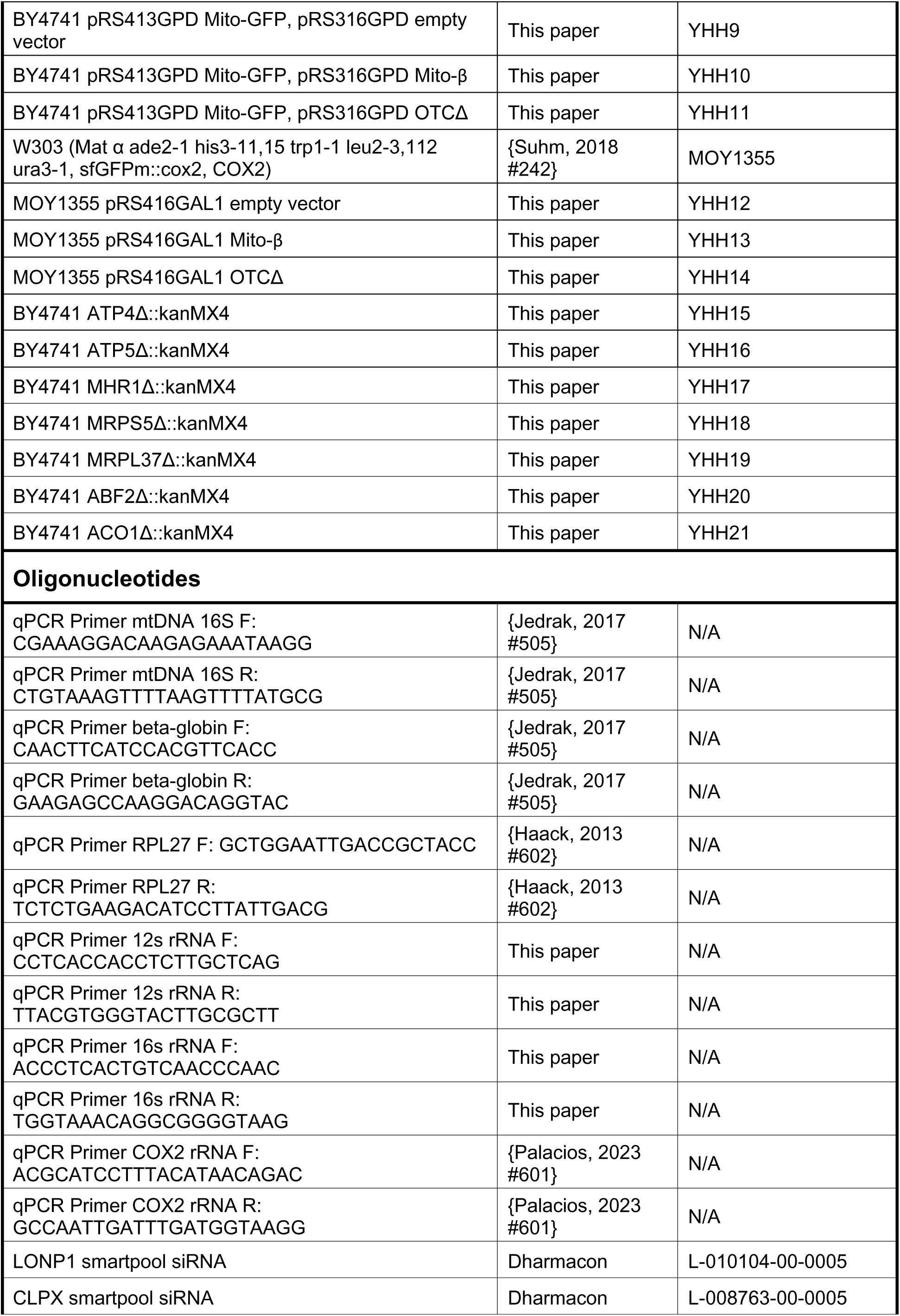
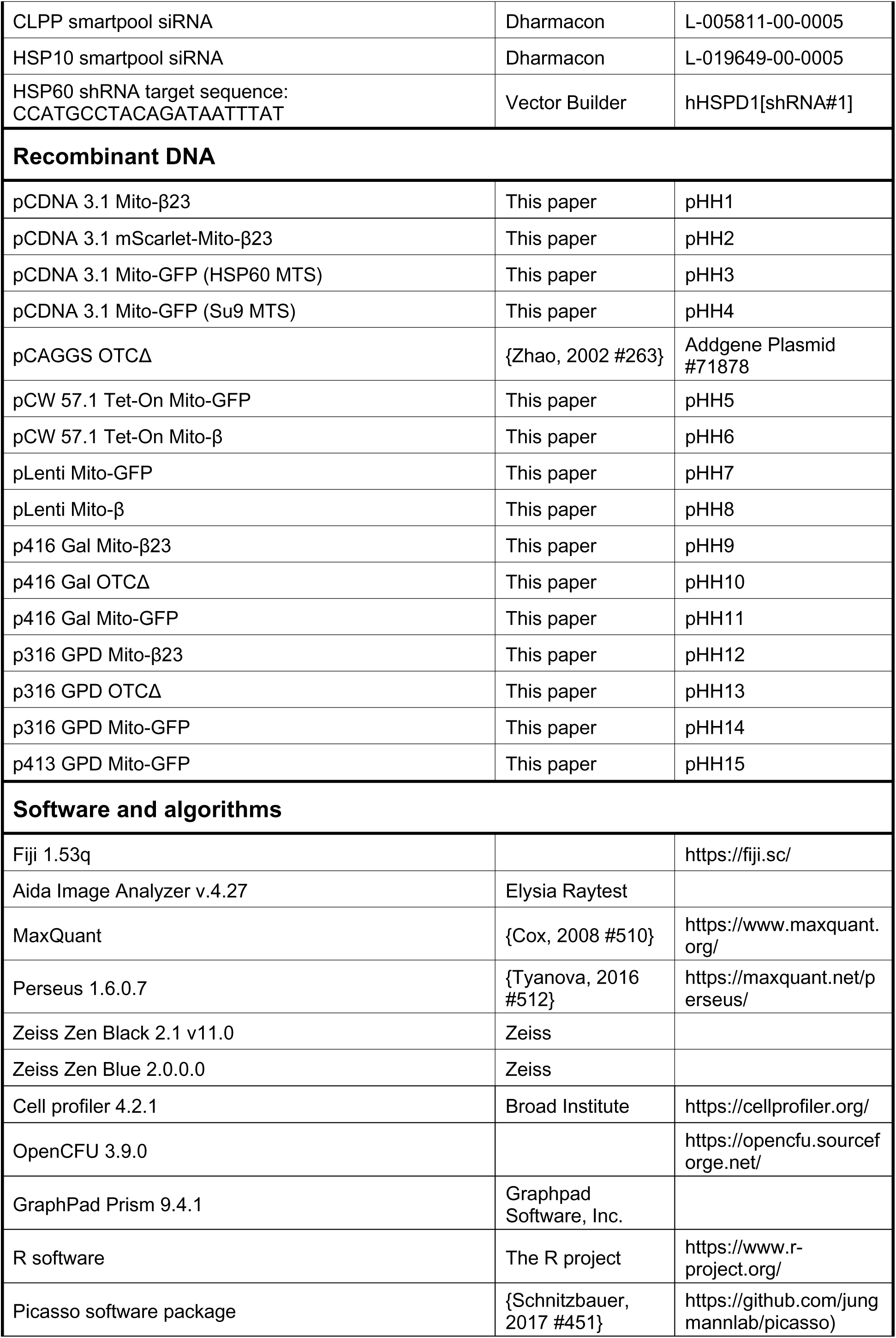
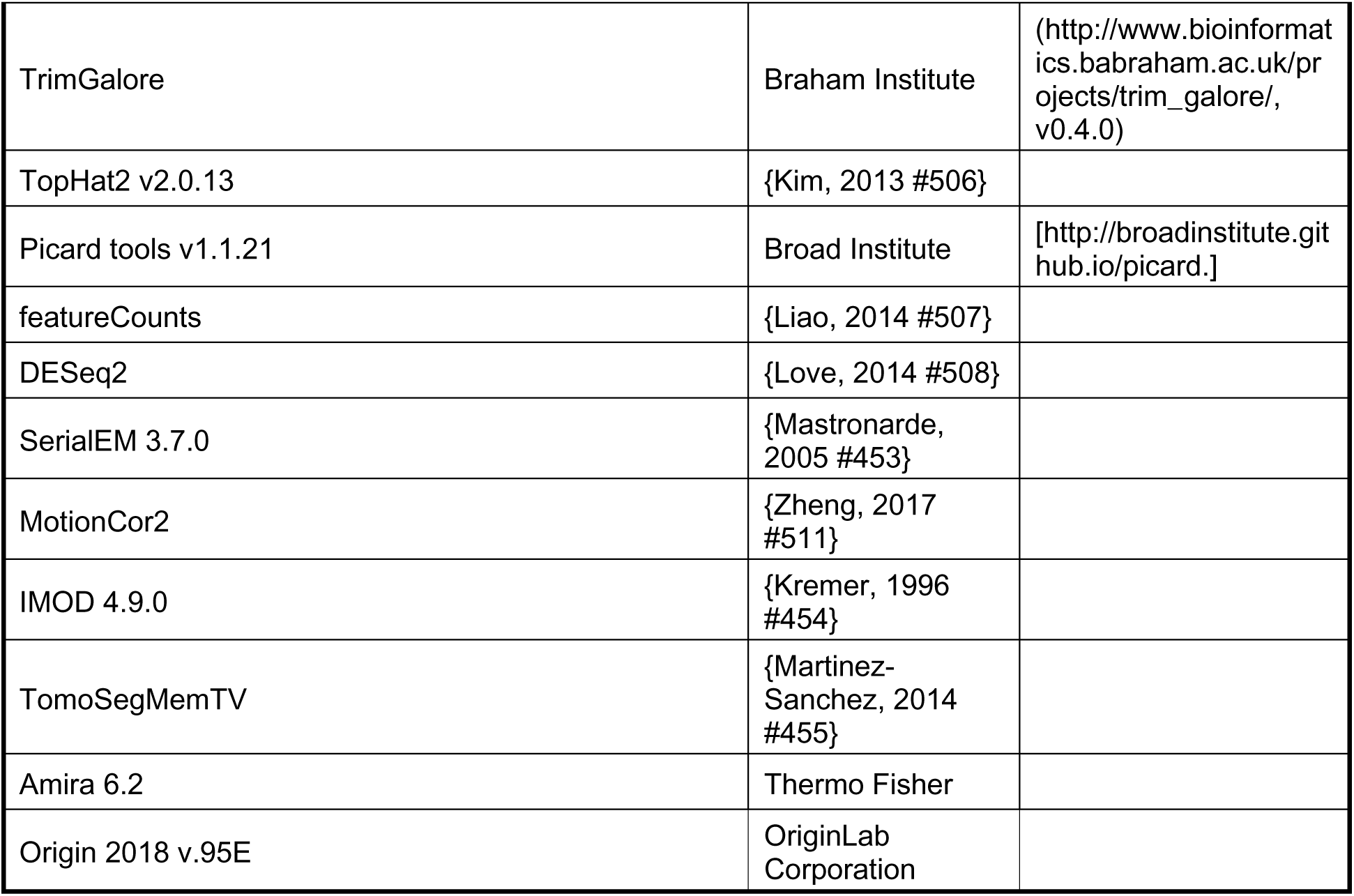

